# Myosin II isoforms play distinct roles in *adherens* junction biogenesis

**DOI:** 10.1101/578997

**Authors:** Mélina L. Heuzé, Gautham Sankara, Tien Dang, Joseph d’Alessandro, Victor Cellerin, David S. Williams, Jan C. M. van Hest, Philippe Marcq, René-Marc Mège, Benoît Ladoux

## Abstract

Adherens junction (AJ) assembly under force is essential for many biological processes like epithelial monolayer bending, collective cell migration, cell extrusion and wound healing. The acto-myosin cytoskeleton acts as a major force-generator during the de novo formation and remodelling of AJ. Here, we investigated the role of myosinII isoforms in epithelial junction assembly. Myosin IIA (NMIIA) and Myosin IIB (NMIIB) differentially regulate biogenesis of adherens junction through association with distinct actin networks. Analysis of junction dynamics, actin organization, and mechanical forces of control and knockdown cells for myosins revealed that NMIIA provides the mechanical tugging force necessary for cell-cell junction reinforcement and maintenance. NMIIB is involved in E-cadherin clustering, maintenance of a branched actin layer connecting E-cadherin complexes and perijunctional actin fibres leading to the building-up of anisotropic stress. These data reveal unanticipated complementary functions of NMIIA and NMIIB in the biogenesis and integrity of AJ.

## Introduction

Tissue integrity and plasticity rely on cell-cell adhesion and cell contractility. The formation, remodelling and disassembly of cell-cell adhesions are fundamental events accompanying all stages of morphogenesis, tissue homeostasis and healing. *Adherens* junctions (AJ) mediated by E-cadherin/catenin complexes are key elements of epithelial cell-cell adhesions and the first ones to assemble upon contact initiation^1–3^. They provide strong mechanical coupling between neighbouring cells through association with the acto-myosin cytoskeleton^4^.

The assembly of *de novo* AJ is crucial for cell-cell rearrangement^5,6^, tissue closure^7^ and the maintenance of epithelial cell integrity during wound healing or cell extrusion^8–10^. During *de novo* cell-cell contacts formation, initial contacts between facing lamellipodia induce immediate clustering of cadherin molecules by trans- and cis-oligomerization^11–14^. Subsequent signalling events involving RhoGTPases trigger local remodelling of the actin cytoskeleton through Arp2/3- or formin-mediated actin polarization in the vicinity of AJs^15–17^. These cytoskeletal rearrangements drive the expansion of cell-cell contacts and inter-cellular adhesion strengthening^2,18,19^.

Non-muscle Myosin II (NMII) has emerged as a fundamental player in force-generation and force-transmission at AJ both *in vitro* and *in vivo*^20–22^. NMII is essential for epithelial tissue architecture^23^, epithelial tissue morphogenesis^24^, tissue repair^25,26^ and cell extrusion^27^. NMII protects junctions from disassembly during development^28^ and provides the mechanical tugging force necessary for AJ reinforcement^29^. In endothelial cells, NMII is recruited early in filopodia-mediated bridge bundles and its activity is required for accumulation of VE-cadherin in nascent AJs^30^. In epithelial cells, NMII favours local concentration of E-cadherin at cell-cell contacts^31,32^ and it is enriched at the edges of elongating junctions where it drives contact expansion in response to RhoA activity^17,18^.

In mammalian cells, NMII heavy chains exist as three different isoforms: NMIIA, NMIIB and NMIIC encoded by *MYH9*, *MYH10* and *MYH14* genes respectively^33,34^. NMIIA and NMIIB are widely expressed whereas NMIIC is not detected in several tissues^35^. Despite structural similarities, NMIIA and NMIIB isoforms have been assigned both redundant and specific functions depending on cell types and processes^36^. NMIIA and NMIIB exhibit different ATPase activities and actin-binding properties ^37–40^, in addition to their specific C-terminal tails that confer them unique functions^41–43^. These two isoforms can exist as activated monomers in cells, but they can also co-assemble as homotypic and heterotypic filaments^44,45^. NMIIA and NMIIB play both unique and overlapping roles *in vivo* ^46–51^. In cells migrating on 2D surfaces, NMIIA localizes at the cell front, limits lamellipodial protrusive activity and reduces 2D cell migration speed by regulating focal adhesions dynamics and traction forces ^52–55^. NMIIB localizes at the cell rear and is required for front-back polarity and tail retraction^53–61^. In 3D, NMIIA favours cell displacement^52–55,62^ while NMIIB drives nuclear translocation^63^. NMIIB also plays a determinant role in durotaxis^64^.

While the roles of NMII isoforms in cell motility on ECM have been extensively studied, very little is known on their respective functions in AJs organization. Smutny and collaborators have reported that NMIIA and NMIIB both localize at apical junction complexes of polarized MCF-7 cells. Upon specific isoform expression silencing, they further proposed that NMIIA may favour the accumulation of E-cadherin in the AJ belt while NMIIB may stabilize the associated perijunctional actin ring^32^. Efimora and collaborators reported an association of NMIIA with contractile actin bundle running parallel to linear AJ in endothelial cells, but failed to precisely localize NMIIB^65^. Here we questioned the unexplored functions of NMII isoforms in epithelial AJ biogenesis using an *in vitro* system based on chemically-switchable micro-patterns, whereby we can control the time and location of a new contact forming between two single cells on a matrix-coated surface.

## Results

### *in vitro* system for the study of early cell-cell contacts

In order to study early AJ biogenesis, pairs of GFP-E-cadherin expressing MDCK cells were plated on arrays of 5 µm-distant fibronectin-coated micro-patterns surrounded by switchable cytorepulsive surfaces^66^. After complete spreading, the confinement imposed by the micro-patterns was released by addition of an RGD-motif containing modified peptide that switched the surface surrounding patterns from a cytorepulsive to an adhesive surface (Supplementary Fig. 1a). Junction biogenesis was monitored by confocal spinning disk microscopy (Fig. 1a, Supplementary Video 1). Within 2 hours, cells extended lamellipodia in random directions and approximately 50 % of the pairs of cells contacted within 12 hours. The junction extended reaching a plateau at 40-45 µm length in around 3 hours (Fig. 1b,d). As previously described^17^, GFP-E-cadherin accumulated at the edges of the junction (Fig. 1b). Once reaching this maximal length, the junction was maintained while showing dynamic retraction-elongation events (Fig. 1b). Importantly, in 98 +/-2 % of the cases, cell-cell contacts were stable and lasted above 3 hours and up to 22 hours (Fig. 1b,e). Analysis of the nucleus-centrosome axis relative to the junction axis showed a relocalization of the centrosome towards the lamellipodia opposite to the cell-cell contact within one hour (Supplementary Fig. 1b,c), as previously reported in different systems and cell types^67–70^. However, although MDCK cells antipolarized in the doublet as if they were initiating a contact inhibition of locomotion, they remained attached to each other in contrast to more mesenchymal cells that proceed with cell separation following repolarization^71^. Together, these observations show that this *in vitro* model system is suitable for the study of early cell-cell contacts at high spatial-temporal resolution.

**Figure 1:**
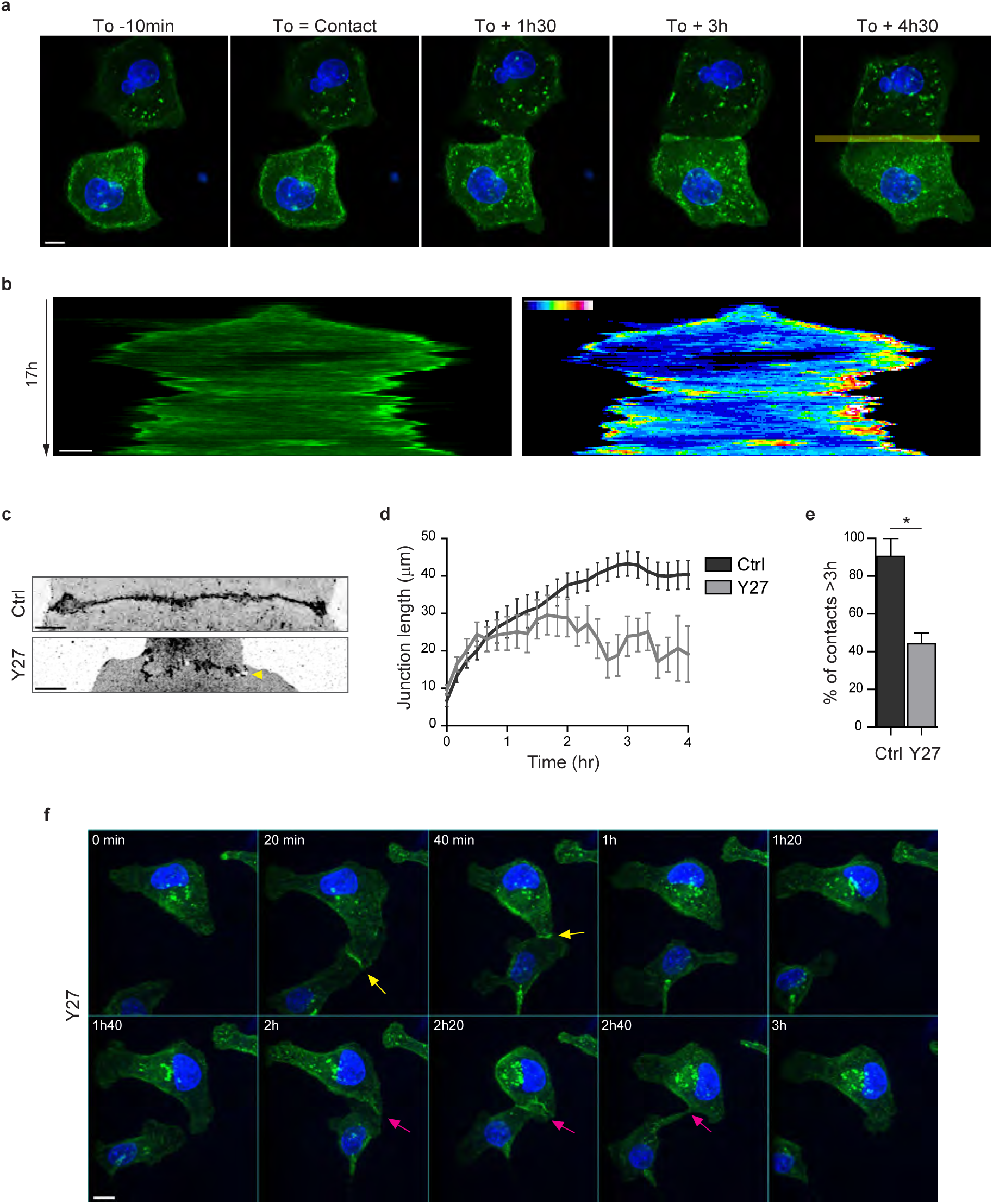
Development of an *in vitro* system for the study of junction biogenesis. **a.** Spinning disk image sequence showing contact extension between two MDCK cells expressing GFP-E-cadherin and stained with Hoechst. Scale bar: 10 µm. **b.** Kymograph of the junction forming in panel b, generated from the yellow line, shown in green and in pseudocolor to highlight GFP-E-cadherin accumulation at junction tips. The junction axis was realigned horizontally for some time points in order to generate the kymograph on a long time scale. Scale bar: 5 µm. **c.** Representative confocal images of β-catenin-stained junctions from MDCK cell doublets. The arrow points at small holes frequently observed within Y27-treated junctions. The cells were fixed 20 hours after addition of BCN-RGD alone or BCN-RGD + Y27 (50 µM). Scale bar: 10 µm. **d.** Graphs showing the evolution of junction length in function of time after contact initiation in Ctrl and Y27-treated MDCK cell doublets. Y27 (50 µM) was added with BCN-RGD. Data are represented as mean +/-SEM. n = 13 and 12 cell doublets from two and three independent experiments, respectively. **e.** Bar graph of the percentage of cell doublets that stay in contact for more than 3 hours in Ctrl and Y27-treated MDCK cells, respectively. Data are represented as mean +/-SEM. n = 13 and 12 cell doublets from two and three independent experiments, respectively. Bonferroni statistical tests were applied for p value. **f.** Spinning disk image sequence of GFP-E-cadherin-expressing MDCK cells pre-stained with Hoechst in the presence of Y27 (50 µM). The sequence starts 3 hours after addition of BCN-RGD + Y27. The arrows highlight transient contacts forming under these conditions. Scale bar: 10 μm.

### NMIIA and NMIIB orchestrate junction biogenesis

To evaluate the involvement of NMII-generated actomyosin contractility in junction biogenesis, we monitored junction formation in cells treated with the ROCK inhibitor Y27632 (Supplementary Video 2). Y27-treated cells exhibited irregular junctions with small digitations and empty spaces and did not elongate as much as in control cells (Fig. 1c,d). They were strongly affected in their capacity to maintain cell-cell contacts, half of the doublets separating before 3 hours (Fig. 1e, f and Supplementary Video 2). Similar results were observed after treating cells with the NMII ATPase activity inhibitor blebbistatin (data not shown) indicating that NMII activity is required for proper junction elongation and stabilization. Furthermore, NMII was required for the centrosome repolarization, as we could not observe any preferential orientation of the nucleus-centrosome axis in Y27-treated doublets (Supplementary Fig. 1d). Next, we explored the involvement of the two NMII isoforms in junction biogenesis. NMIIA has been reported to be by large the major isoform of NMII expressed in MDCK cells^35^. However, immunostainings revealed that the three isoforms, NMIIA, NMIIB and NMIIC could be detected in MDCK cells. NMIIA and NMIIC fully co-localized to similar structures, which was not the case for NMIIB (Supplementary Fig. 3a,b). For these reasons, we decided to focus on NMIIA and NMIIB isoforms. Expression of each isoform was silenced in GFP-E-cadherin MDCK cells by stable transfection of specific ShRNA encoding plasmids, leading to an inhibition of expression of around 60-70% (Fig. 2a,b and Supplementary Fig. 2a,b). The analysis of cell-cell contact formation in cell doublets by live-imaging (Supplementary Video 3) revealed that NMIIB knock-down (NMIIB KD) cells formed and extended intercellular junctions very similar to control (Ctrl) cells (Fig. 2c-f). In contrast, almost half of NMIIA knock-down (NMIIA KD) cell doublets were unable to sustain contacts more than 3 hours, and when they did so, these contacts remained shorter than for Ctrl or NMIIB KD cell doublets (Fig. 2c-f), similar to what was observed in Y27-treated cell doublets. NMIIB KD doublets, despite their ability to maintain cell-cell contacts for longer times, formed twisted junctions that were significantly less straight than Ctrl and NMIIA KD cells and deviated significantly more from their initial orientation (Fig. 2g,h). These defects in NMIIB KD cells were already observed at early stages of junction biogenesis and were associated to the formation of large extensions of junctional membrane (Fig.2i, arrows). Together, these results show that both NMIIA and NMIIB are required for the biogenesis of stable AJs, albeit with different contributions; NMIIA favours temporal stability whereas NMIIB ensures the straightness and spatial stability of the junctions.

**Figure 2:**
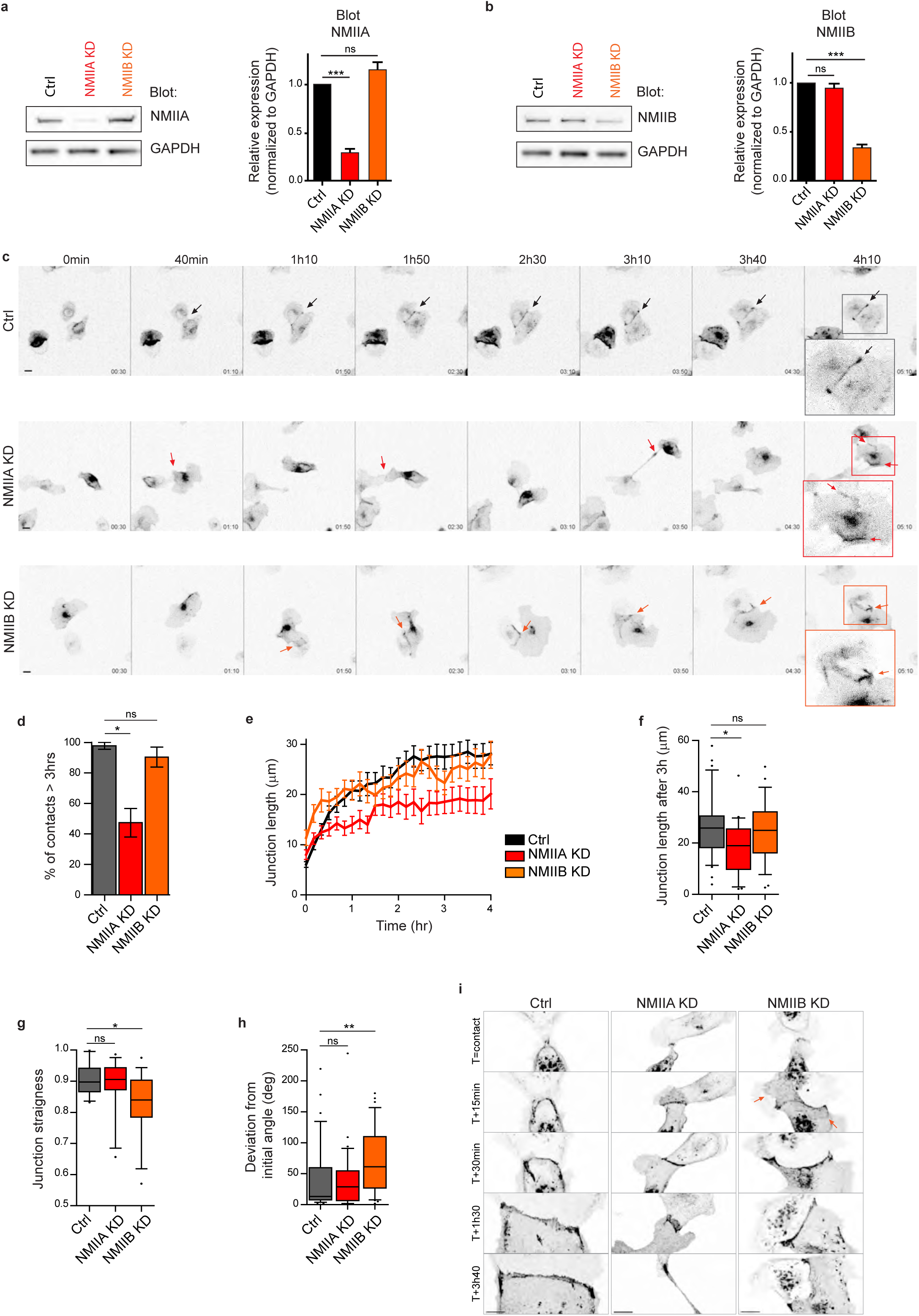
NMIIA and NMIIB are both required for proper junction biogenesis. **a.b.** Left panels: Representative immunoblots showing the isoform specific knockdown of NMIIA (a) and NMIIB (b) in NMIIA KD and NMIIB KD MDCK cells. GAPDH expression levels were used as loading controls. Right panels: Bar graphs showing the relative expression level of NMIIA and NMIIB proteins in Ctrl, NMIIA KD and NMIIB KD cells normalized to GAPDH expression levels. Data are represented as mean +/-SEM from three independent experiments. Kruskall-Wallis statistical tests were applied for p value. **c.** Representative epifluorescent image sequences of GFP-E-cadherin over a time course of 5 hours showing the dynamics of junction formation at low magnification in Ctrl, NMIIA KD and NMIIB KD MDCK cells. The arrows indicate the position and the orientation of the junctions. Scale bar: 10 µm. **d.** Bar graph of the percentage of cell doublets that stay in contact for more than 3 hours. Data are represented as mean +/-SEM. Tukey’s multiple comparison statistical tests were applied for p value. n = 36, 37 and 31 cell doublets for Ctrl, NMIIA KD and NMIIB KD cells respectively, from three independent experiments. **e.** Plots showing the evolution of junction length in function of time for Ctrl, NMIIA KD and NMIIB KD cell doublets. Data are represented as mean +/-SEM. n = 40, 43 and 35 cell doublets for Ctrl, NMIIA KD and NMIIB KD cells respectively, from four independent experiments. **f.** Box & whiskers graphs representing the junction length after 3 hours after contact, for Ctrl, NMIIA KD and NMIIB KD cell doublets. n = 34, 21 and 28 cell doublets for Ctrl, NMIIA KD and NMIIB KD cells respectively, from four independent experiments. **g.** Box & whiskers graphs showing the junction straightness (calculated as the euclidean/accumulated length ratio) in Ctrl, NMIIA KD and NMIIB KD cell doublets 2 hours after contact. n = 12, 15 and 17 cell doublets for Ctrl, NMIIA KD and NMIIB KD cells respectively, from three independent experiments. **h.** Box & whiskers graph showing the angular deviation of junctions during the 3 first hours of contact in Ctrl, NMIIA KD and NMIIB KD cell doublets. n = 35, 30 and 32 cell doublets for Ctrl, NMIIA KD and NMIIB KD cells respectively, from four independent experiments. **f-h:** Mann-Whitney statistical tests were applied for p value. **i.** Representative spinning disk GFP-E-cadherin image sequences over a time course of 4 hours showing the dynamics of junction formation at high magnification in Ctrl, NMIIA KD and NMIIB KD MDCK cells. The red arrows point at junctional extensions typically observed in NMIIB KD doublets. Scale bar: 10 µm.

### NMIIB localizes to a junctional actin pool distinct from perijunctional NMIIA-associated contractile fibres

To better understand the respective roles of NMIIA and NMIIB in junction biogenesis, we next studied their subcellular localization at nascent cell-cell contacts in cell doublets. Immunostainings revealed that the localization of the two isoforms was mutually exclusive. Anti-NMIIA antibodies stained actin bundles that were parallel to the junction but set at 1 to 2 μm from it. NMIIA was also found associated to actin cables parallel to the cortex of non-junctional membranes (Fig. 3a,b,e and Supplementary Fig. 3a) in addition its association to the classical ventral stress fibres. In contrast, NMIIB immunostaining was associated with the junctional plasma membranes as well as with a cytoplasmic network (Fig. 3c-e and Supplementary Fig. 3b), that was identified as the vimentin intermediate filament network as reported by Menko and colleagues^72^ in lens epithelial cells. Importantly, the localization of each isoform was not affected by the silencing of the other isoform (Supplementary Fig.3c). Smutny and colleagues previously reported that NMIIA and NMIIB both localized to apical epithelial junctions in polarized MCF-7 cells^32^. Given that, we followed the localization of both isoforms during apico-basal polarization of MDCK cells (Fig. 3f and Supplementary Fig. 3d). After 3 days of culture, confluent MDCK cells started to develop an apico-basal polarization and the two isoforms colocalized to apically positioned *zonulae adherens* colocalizing with E-cadherin as previously reported in MCF-7 cells^32^. At the ventral side, they were associated to stress fibres. However, we confirmed a different localization of NMIIB and NMIIA in sub-confluent cell clusters after one day of culture. NMIIA was still associated to stress fibres while NMIIB colocalized with E-cadherin at cell-cell contacts. These differential distributions at the early stages of AJ formation were not specific to MDCK cells, and were observed as well in small clusters of Caco2 cells (Supplementary Fig.3e). Considering recent findings showing a possible interaction between NMIIB and α-catenin^73^, we hypothesized that NMIIB could be recruited to the junction through α-catenin/E-cadherin complexes. Accordingly, in α-catenin KD MDCK cells^74^, NMIIB was relocalized to NMIIA-enriched stress fibres and circumnuclear actin cables (Fig. 3e), indicating that α-catenin is required for NMIIB junctional recruitment.

**Figure 3:**
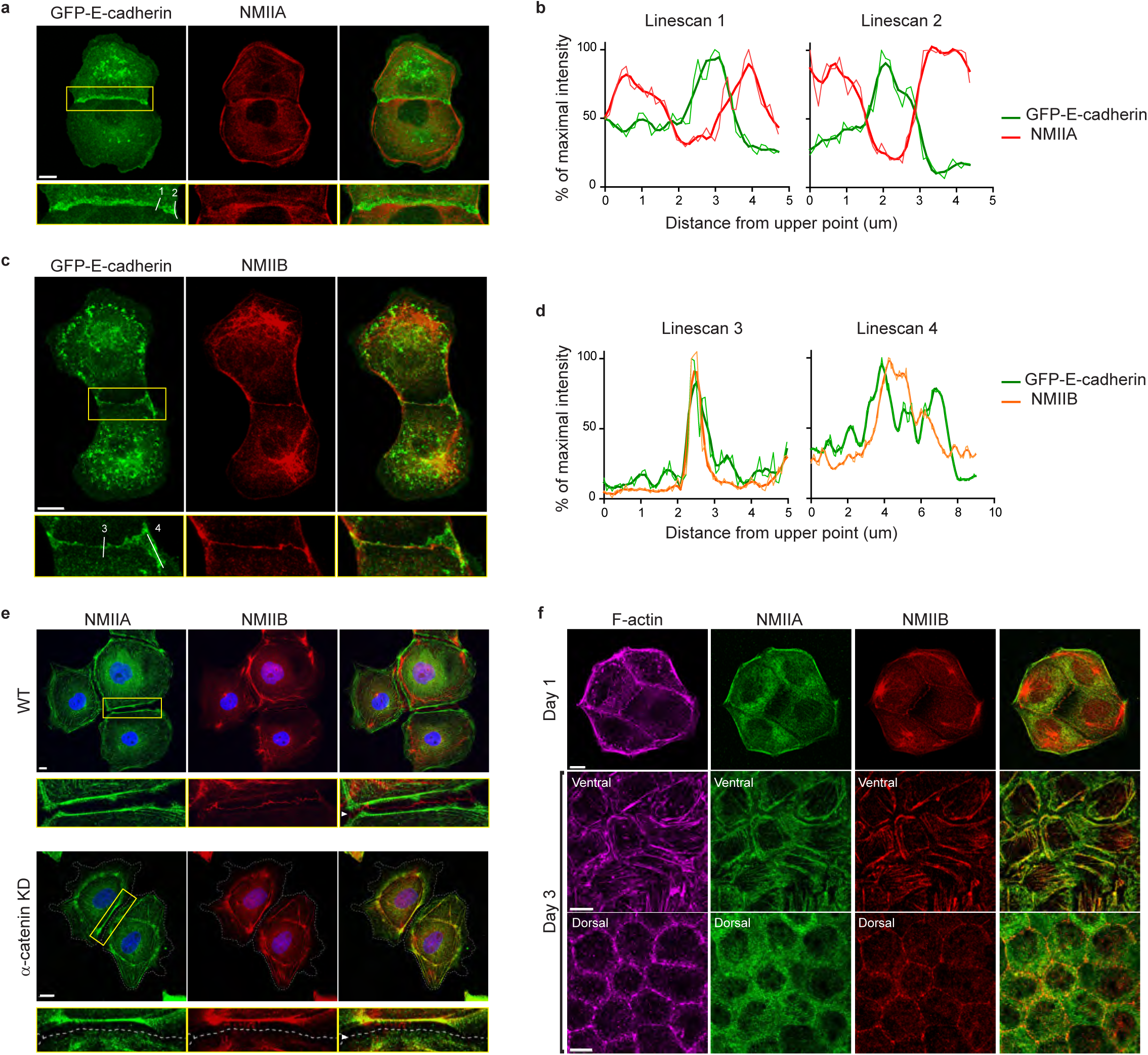
NMIIB, but not NMIIA, localizes to early AJs. **a.c.** Representative confocal images and zoom boxes of GFP-E-cadherin-expressing MDCK cell doublets fixed 20 h after BCN-RGD addition and immuno-stained for NMIIA (a) or NMIIB (c). Scale bar: 10 µm. **b.d.** Relative intensity profiles (raw and smoothed data) of GFP-E-cadherin and NMIIA (b) or NMIIB (d) signals along the lines represented in (a) and (c) respectively. **e.** Representative confocal images and zoom boxes of WT (upper panel) or α-catenin KD (lower panel) MDCK cells fixed 20 h after BCN-RGD addition and immuno-stained for NMIIA and NMIIB. White arrow heads indicate the cell-cell contact which is depicted as a dotted line in α-catenin KD MDCK cells. Scale bar: 10 µm. **f.** Representative confocal images of WT MDCK cells plated on fibronectin-coated glass for 1 or 3 days and stained for F-actin, NMIIA and NMIIB. Scale bar: 10 µm.

To better characterize the organization of the actomyosin cytoskeleton at nascent AJs, co-stainings of NMIIA, NMIIB, F-actin and β-catenin performed on control MDCK cells were imaged using structured illumination microscopy (SIM). NMIIA was associated to thick F-actin bundles running parallel to, and located a few microns away from the junctional membranes (Fig. 4a), as reported for NMIIA localization in linear junctions of endothelial cells^30,65^. We confirmed at this resolution that NMIIA did not colocalize with β-catenin-labelled cadherin-catenin complexes. Interestingly, NMIIA appeared distributed on these actomyosin bundles in sarcomere like structures as described before in other cellular contexts^75,76^. These NMIIA labelled structures were almost free of NMIIB staining. NMIIB junctional staining colocalizing with β-catenin was associated with a 200 nm to 1 µm thick fuzzy F-actin network (Fig. 4a,b). This junctional F-actin network was labelled by both Arp2/3 and cortactin (Fig. 5b,c) thus corresponding to an Arp2/3-nucleated branched actin meshwork. Looking at short junctions that probably corresponded to nascent cell-cell contacts, we could also observe the strong enrichment of NMIIB and the exclusion of NMIIA at the contact zone (Fig.4c,d).

**Figure 4:**
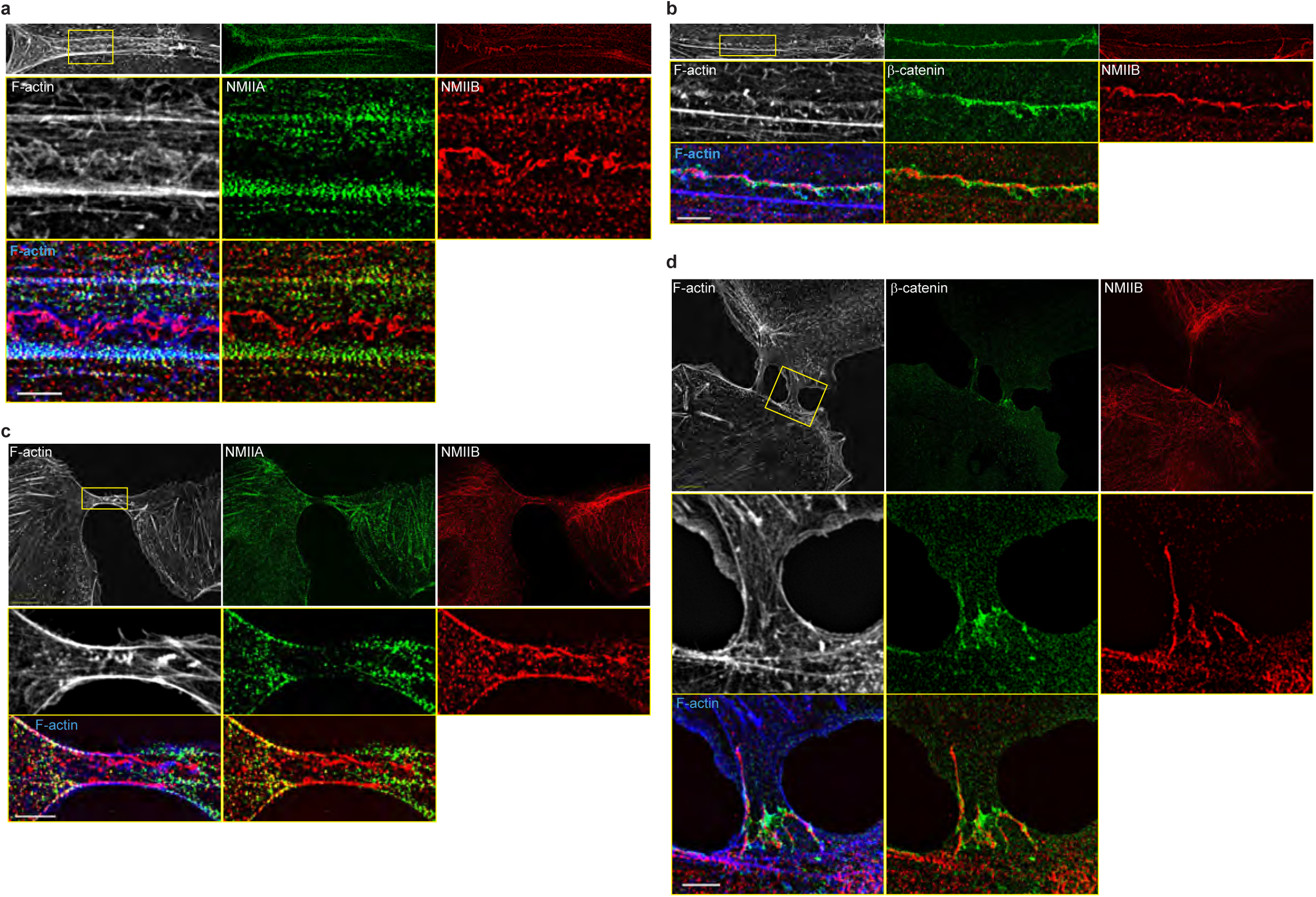
NMIIB localizes to a junctional actin network distinct from NMIIA-associated actin. **a-d.** SIM (Structured Illumination Microscopy) images of WT MDCK cells fixed 20h after addition of BCN-RGD and stained as indicated. Scale bar: 3 µm.

**Figure 5:**
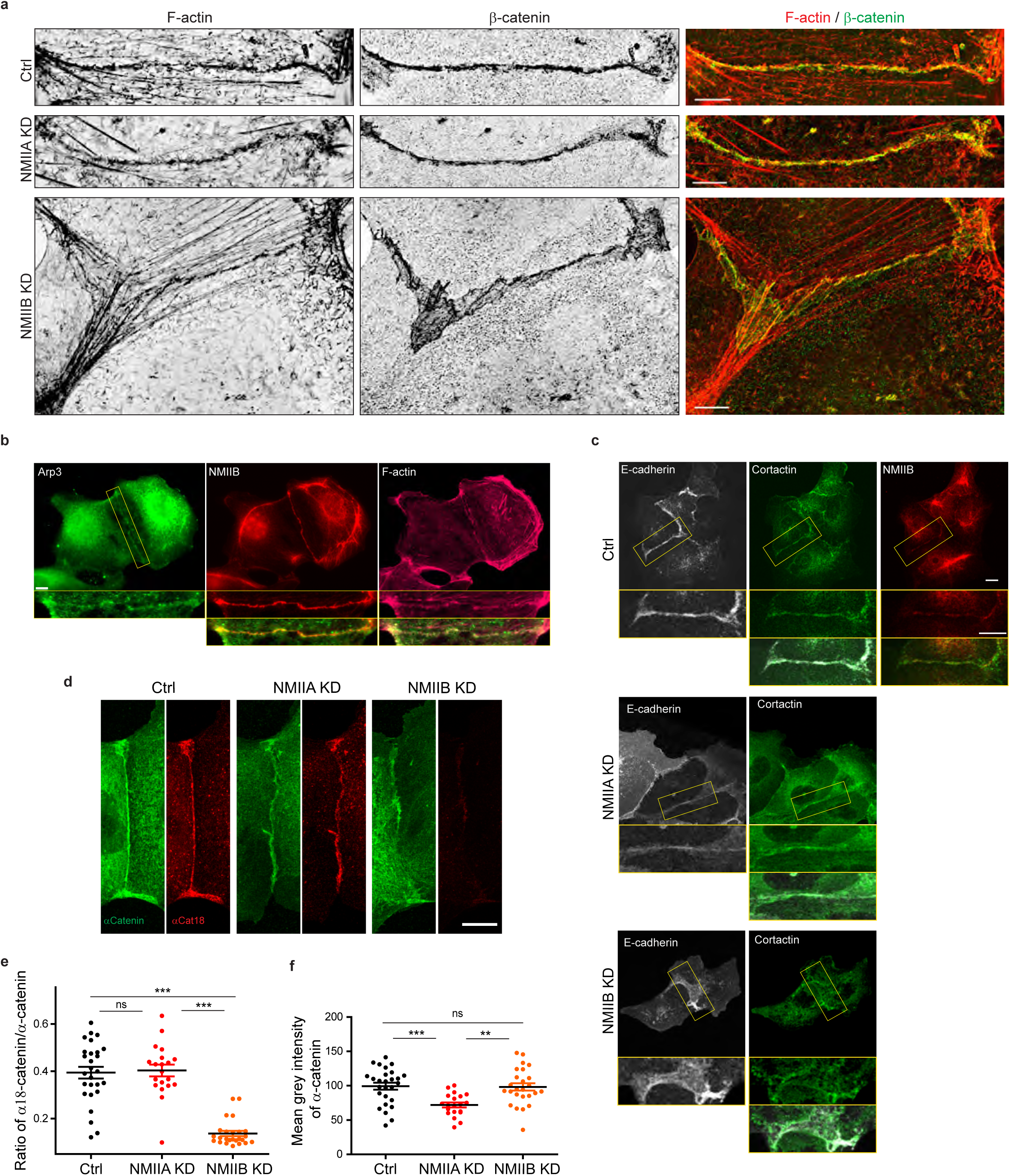
NMIIB supports junctional branched actin organization and regulates α-catenin unfolding. **a.** SIM (Structured Illumination Microscopy) images of junctional areas from Ctrl, NMIIA KD and NMIIB KD cells fixed 20 h after addition of BCN-RGD and stained for F-actin and β-catenin. Scale bar: 5 µm. **b.c.** Representative epifluorescent (b) or confocal (c) images with zoom boxes of MDCK cells and stained as indicated. Scale bar: 10 µm. **d.** Representative confocal images of junctional area from Ctrl, NMIIA KD and NMIIB KD cells stained for α-catenin and α-cat18. Scale bar: 10 µm. **e.f.** Scatter plots with mean +/-SEM showing the ratio of junctional α-cat18/α-catenin signals (e) and the mean intensity levels of α-catenin signal at the junction (f). n = 27, 20, 25 cell doublets for ctrl, NMIIAKD and NMIIBKD, respectively from two independent experiments. Kruskal-Wallis statistical tests were applied for p value.

Altogether, these observations reveal that early during AJ biogenesis, NMIIB is exclusively associated to a junctional Arp2/3-nucleated F-actin, structurally distinct from the perijunctional contractile NMIIA-associated actin bundles running parallel to the junction.

### NMIIA regulates the organization of perijunctional actin bundles while NMIIB supports the maintenance of the junctional branched actin layer

Based on these observations, we subsequently explored the possibility that NMIIB and NMIIA could differentially regulate actin assembly at the junction, thereby maintaining its structural integrity. Using SIM microscopy, we analyzed the organization of junctional actin cytoskeleton in NMIIA KD and NMIIB KD cells. NMIIA KD cells exhibited shorter actin bundles running parallel to the junction, while their junctional F-actin meshwork was comparable to the one of Ctrl cells, both in terms of morphology and cortactin staining (Fig. 5b,c and Supplementary Fig. 4). In contrast, NMIIB KD cells presented a strongly enlarged area of junctional F-actin meshwork colocalizing with β-catenin that corresponded to overlapping membrane extensions stained with cortactin (Fig.5b,c and Supplementary Fig.4). In addition, while they retained some of the perijunctional actin bundles, we could observe numerous oblique actin bundles directed toward the junction (Fig.5c and Supplementary Fig.4). These results show that NMIIA supports the organization of perijunctional actin bundles while NMIIB is required for the regulated assembly of junctional branched F-actin that couples perijunctional bundles to the plasma membrane.

The Arp2/3-nucleated actin network at the *zonula adherens* has been shown to regulate junctional tension in epithelial monolayers ^77^. On the other hand, junctional tension has been shown to associate with the presence of α-catenin molecules under open conformations ^78,79^. Moreover, a direct link between α-catenin and NMIIB has been reported^73^, suggesting that NMIIB recruitment, α-catenin molecular unfolding and regulation of Arp2/3-dependent branched actin polymerization could be tightly linked. Thus, we performed immunostainings with the α18 monoclonal antibody recognizing the open conformation of the protein^79^. Strikingly, the ratio of α18 on total α-catenin junctional staining was decreased by four times in NMIIB KD cells compared to Ctrl cells, while it was not affected in NMIIA KD cells. This suggests that junctional α-catenin molecules were significantly turned to the closed conformational state in NMIIB KD cells (Fig.5d-f). In contrast, the total α-catenin junctional levels were significantly reduced in NMIIA KD cells, as shown by others^31,32^. Taken together, these results strengthen complementary contributions for NMIIB and NMIIA where NMIIB is the main isoform required for the organization of junctional branched actin and NMIIA for organization of perijunctional contractile actin fibres.

### NMIIA is required for the generation of forces at E-cadherin adhesions while NMIIB favours their transmission through F-actin anchoring

The formation of cell-cell junctions in cell doublets is concomitant with the formation of cell-matrix adhesions and the tugging force applied on cell-cell contacts must be compensated by traction of the cells on cell-matrix adhesion complexes^29,80–82^. To further understand the contributions of NMII isoforms in junction biogenesis, we thus experimentally decoupled these two adhesion systems. We first investigated the role of NMII isoforms in cell-matrix adhesion by seeding single Ctrl, NMIIA KD and NMIIB KD cells on fibronectin-coated glass. NMIIA KD cells spread 1.7 times more than Ctrl and NMIIB KD cells on fibronectin and their actin cytoskeleton was highly perturbed exhibiting a strong decrease in ventral stress fibres and cortical actin bundles together with an enlargement of their lamellipodia (Fig. 6a,b). NMIIA KD cells also formed significantly less focal adhesions (Fig. 6a,c). In contrast, NMIIB KD cells showed no defect in actin organization, cell spreading or focal adhesion formation (Fig. 6a-c). Next, we measured by TFM the magnitude of traction forces applied by single cells on deformable fibronectin-coated 30 kPa PDMS gels. NMIIA KD cells exerted lower traction forces than Ctrl cells as reported by others^55,61^. NMIIB KD cells, on the contrary, did not show any defect in traction force generation on this substratum (Fig. 6d,e). These results, in agreement with previous studies^55,57^, show that NMIIA is the isoform regulating cell spreading, cell adhesion, traction force generation and organization of contractile actin structures on fibronectin. In contrary MNIIB is not contributing at all to the cell-matrix adhesion, focal adhesion formation, actomyosin reorganization and traction forces on fibronectin.

**Figure 6:**
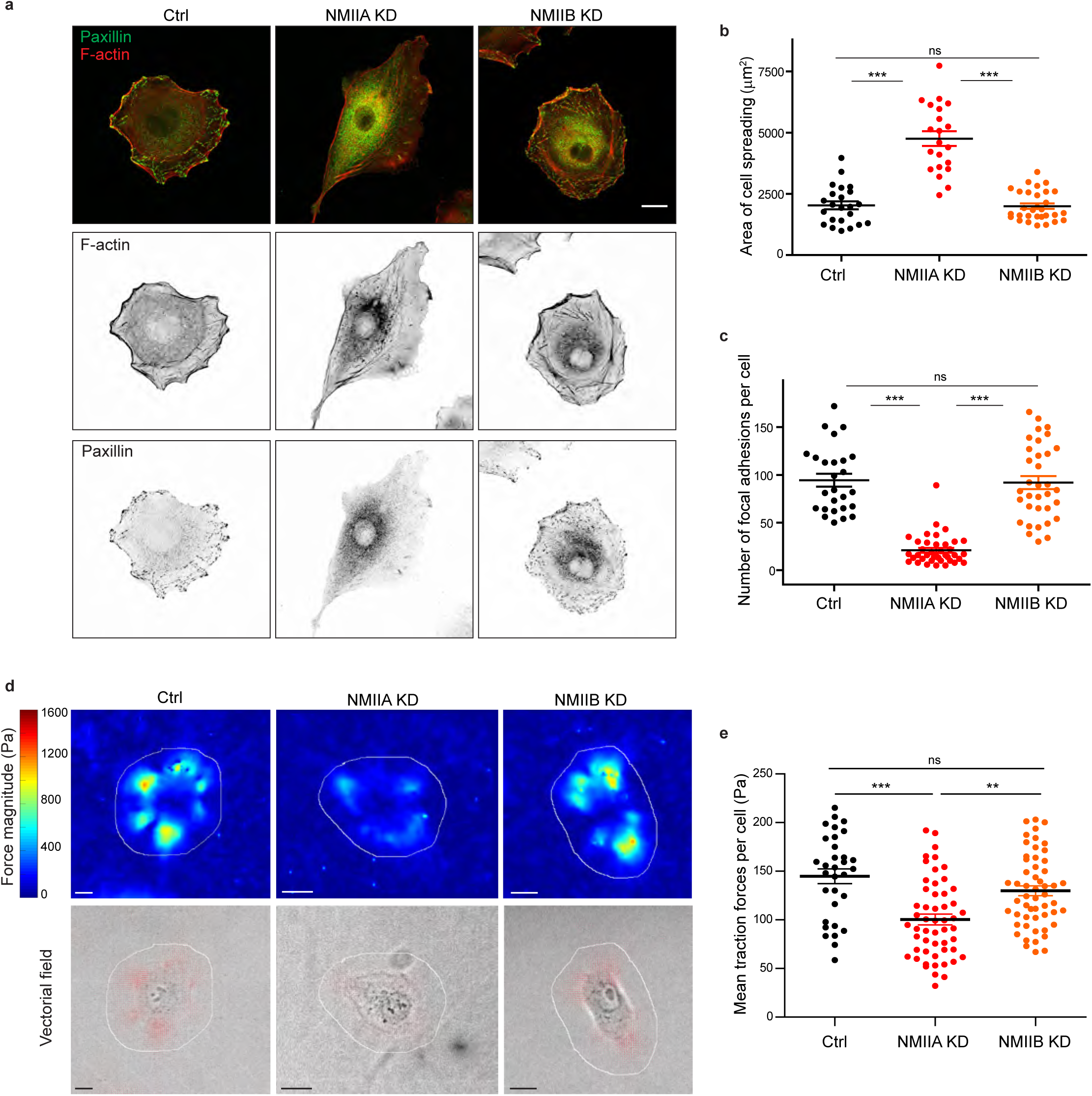
NMIIA, but not NMIIB, regulates cell-matrix adhesions and traction forces. **a.** Representative confocal images of paxillin and F-actin staining of Ctrl, NMIIA KD and NMIIB KD single cells plated on fibronectin-coated glass coverslip for 16 hours. Scale bar: 10 µm. **b.c.** Scatter plots with mean +/-SEM showing the spreading area (b) and number of focal adhesions (c) of Ctrl, NMIIA KD and NMIIB KD single cells plated on fibronectin for 16 hours. n = 23, 21 and 30 cells for (b) and 26, 39 and 34 cells for (c) respectively, from two independent experiments. Kruskal-Wallis statistical tests were applied for p value. **d.** Heat map (upper panel) and vectorial field (lower panel) representing respectively the magnitude and the orientation of traction forces exerted by the single Ctrl, NMIIA KD and NMIIB KD cells, on fibronectin-coated PDMS deformable substrate (30 KPa). Cell masks used for quantification are drawn in white. Scale bar: 10 µm. **e.** Scatter plots with mean +/-SEM showing the mean traction forces exerted by single Ctrl, NMIIA KD and NMIIB KD cells. n = 32, 53 and 54 cells, respectively. Kruskall-Wallis statistical tests were applied for p value.

To explore the contribution of NMII isoforms to E-cadherin-mediated cell-cell adhesion per se, we seeded single cells on E-cadherin-coated substrates (Fig. 7a,b). After 6 hours, Ctrl and NMIIA KD cells had spread similarly with mean areas of 1178 ± 40 µm² and 1031 ± 37 µm^2^ respectively, while NMIIB KD cells spreading was significantly reduced (mean area = 515 ± 21 µm^2^) (Fig. 7a,c). Ctrl cells organized thick circumnuclear actin arcs, as well as radial actin fibres connected to peripheral β-catenin clusters (Fig. 7a), as previously described^83,84^. NMIIA KD cells, while spreading as Ctrl cells on E-cadherin and forming cadherin clusters in similar numbers, lacked the circumnuclear actin arcs (Fig. 7a,c,d). In contrast, NMIIB KD cells kept the organization of circumnuclear actin arcs, but were depleted of radial actin bundles, did not form significant β-catenin clusters and failed to spread on E-cadherin (Fig. 7a,c,d). This data indicated that NMIIB, but not NMIIA, plays a major role in the clustering and stabilization of E-cadherin/catenin complexes that in turn promote cell spreading. Our findings also suggest that NMIIA, but not NMIIB, is required for the formation of contractile actin fibres that apply traction forces on the cadherin adhesions. We thus measured the capacity of these cells to transmit forces through E-cadherin complexes by TFM, seeding them on E-cadherin-coated 15 kPa PDMS elastic gels. Compared to Ctrl cells, NMIIA KD cells exhibited very low forces on E-cadherin substrate (Fig.7e,f), confirming that NMIIA generates the forces transmitted to E-cadherin adhesions. NMIIB KD cells, that failed to cluster cadherin/catenin complexes, also generated lower traction forces than Ctrl cells, albeit to a lesser extent than NMIIA KD cells (Fig. 7e, f). Even though both myosin isoforms contribute to cell-generated forces on E-cadherin substratum, they have complementary contributions. NMIIA is required for the formation of stress fibres while NMIIB would rather regulate the transmission of force and the coupling of actin stress fibres to the cadherin-catenin complexes.

**Figure 7:**
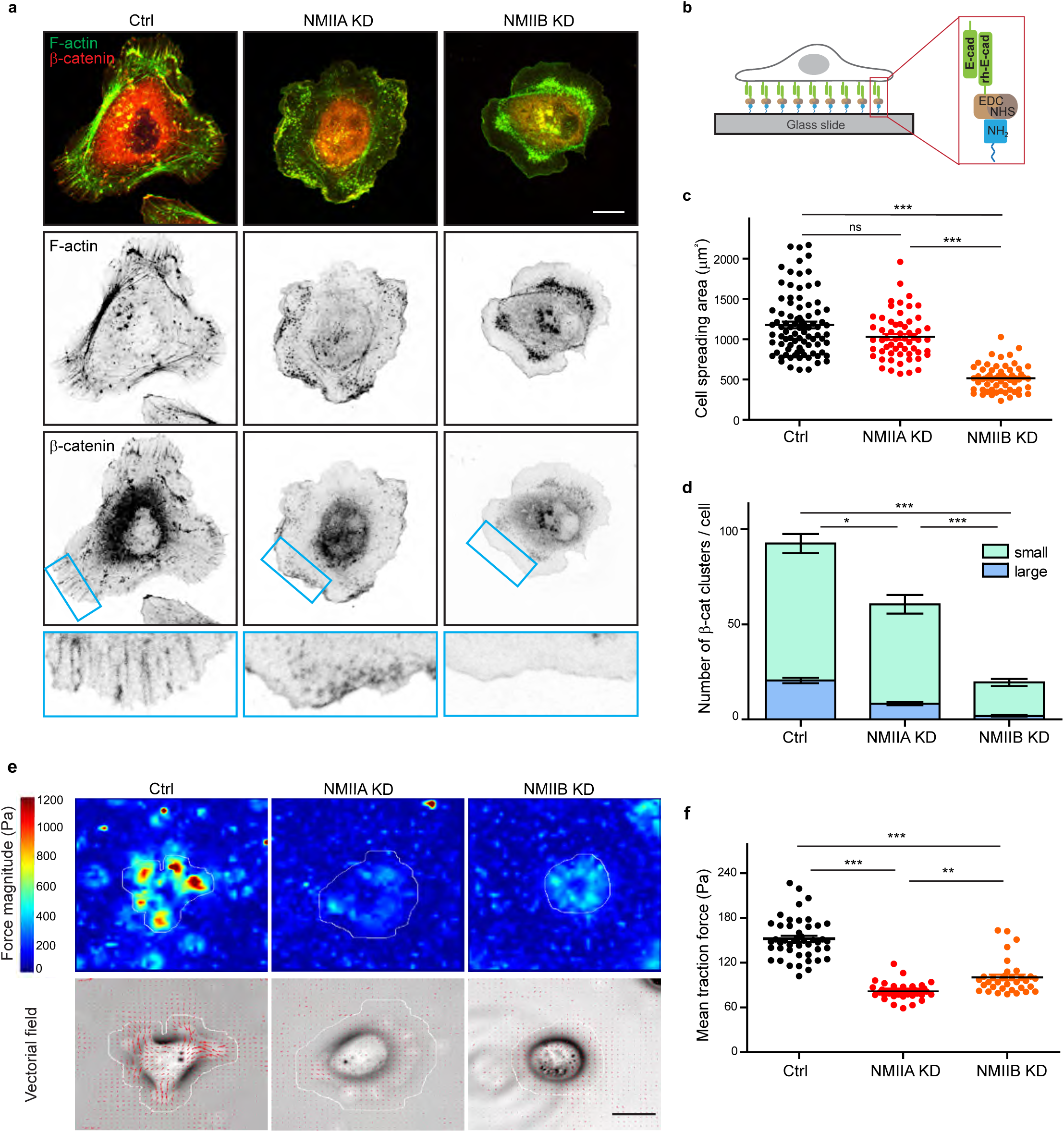
NMIIB favours E-cadherin clustering on E-cadherin-coated substrate. **a.** Confocal images with zoom boxes of Ctrl, NMIIA KD and NMIIB KD cells plated on E-cadherin-coated glass for 6 hours and immuno-stained for β-catenin and F-actin. Scale bar: 10 µm. **b.** Scheme depicting the experimental set-up. **c.** Scatter plots with mean +/-SEM showing the cell spreading area of Ctrl, NMIIA KD and NMIIB KD cells plated on E-cadherin coated glass after 6 hours. n = 87, 58 and 58 cells respectively from two independent experiments. Kruskal-Wallis statistical tests were applied for p value. **d.** Bar graph showing the number of β-catenin clusters per cell in Ctrl, NMIIA KD and NMIIB KD cells plated on E-cadherin coated glass. The clusters were classified in two categories: large clusters with area larger than 1 µm², and small clusters with area ranging from 0.2 µm² to 1 µm². Data are represented as mean +/-SEM, n = 26, 27 and 26 cells respectively from two independent experiments. Kruskal-Wallis statistical test were applied for p value **e.** Heat map (top panel) and vectorial field (bottom panel) representing respectively the magnitude and the orientation of traction forces exerted by the single Ctrl, NMIIA KD and NMIIB KD cells on E-cadherin coated PDMS deformable gels (15 KPa). Cell masks used for quantification are drawn in white. Scale bar: 20 µm. **f.** Scatter plots with mean +/-SEM showing the mean traction forces exerted by Ctrl, NMIIA KD and NMIIB KD cells on E-cadherin coated PDMS deformable gels (15 KPa). n = 46, 34 and 34 cells respectively from two independent experiments. Kruskal-Wallis statistical tests were applied for p value.

### NMIIA and NMIIB are required for proper organization of inter-cellular junctional stress

To directly determine how NMIIA and NMIIB contribute to traction force generation and transmission during AJ biogenesis, we mapped traction forces before and after cell-cell contact formation in cell doublets. Hotspots of traction forces were generated at the periphery of the doublet where lamellipodia arise (Fig.8a). As expected from the TFM data obtained with single cells seeded on fibronectin, NMIIA KD doublets, compared to Ctrl and NMIIB KD ones, exhibited very low traction forces both before and after cell-cell contact formation (Fig. 8a-c). NMIIB KD doublets developed traction forces similar in magnitude to those developed by Ctrl ones, with however different patterns. Hotspots of forces frequently appeared in the junctional area in NMIIB KD doublets that were generally absent in Ctrl and NMIIA doublets (Fig. 8a). We quantified these differences by analysing the spatial repartition of forces in the peripheral and central subdomains of the junction, and their orientation relative to main junction axis (parallel, F_//_, and perpendicular, F_⊥_, components). NMIIB KD doublets generated higher F_⊥_ in the central part of the junction and lower values of F_//_(albeit not significantly) with respect to Ctrl doublets in both the peripheral and the central part of the junction (Supplementary Fig.5a-c). These results show that NMIIB plays an important role in the repartition of traction forces under the junction and that NMIIA is essential for the generation of traction forces in general. We next quantified the capacity of NMIIA KD and NMIIB KD cells to transmit forces across the junction. Following Newton’s laws, the net traction force exerted by an isolated doublet is zero, up to the measurement noise. Conversely, the net traction forces exerted by each of the two cells are equal in magnitude and opposite in direction, compensating exactly^29,80,81^. We thus calculated the resultant vectorial sum of forces per cell (Fig. 8b). In all conditions, the resultant force per cell before contact was within the level of noise as expected for isolated cells and increased within 30 minutes after contact to reach a plateau, attesting the capacity of all three cell lines to transmit intercellular tugging forces across the junction (Fig.8b,c). However, in NMIIA KD cells, the resultant forces per cell at the plateau was significantly lower than in Ctrl and NMIIB KD cell doublets (Fig.8c), which is consistent with the inability of these cells to apply strong traction forces on fibronectin substratum.

**Figure 8:**
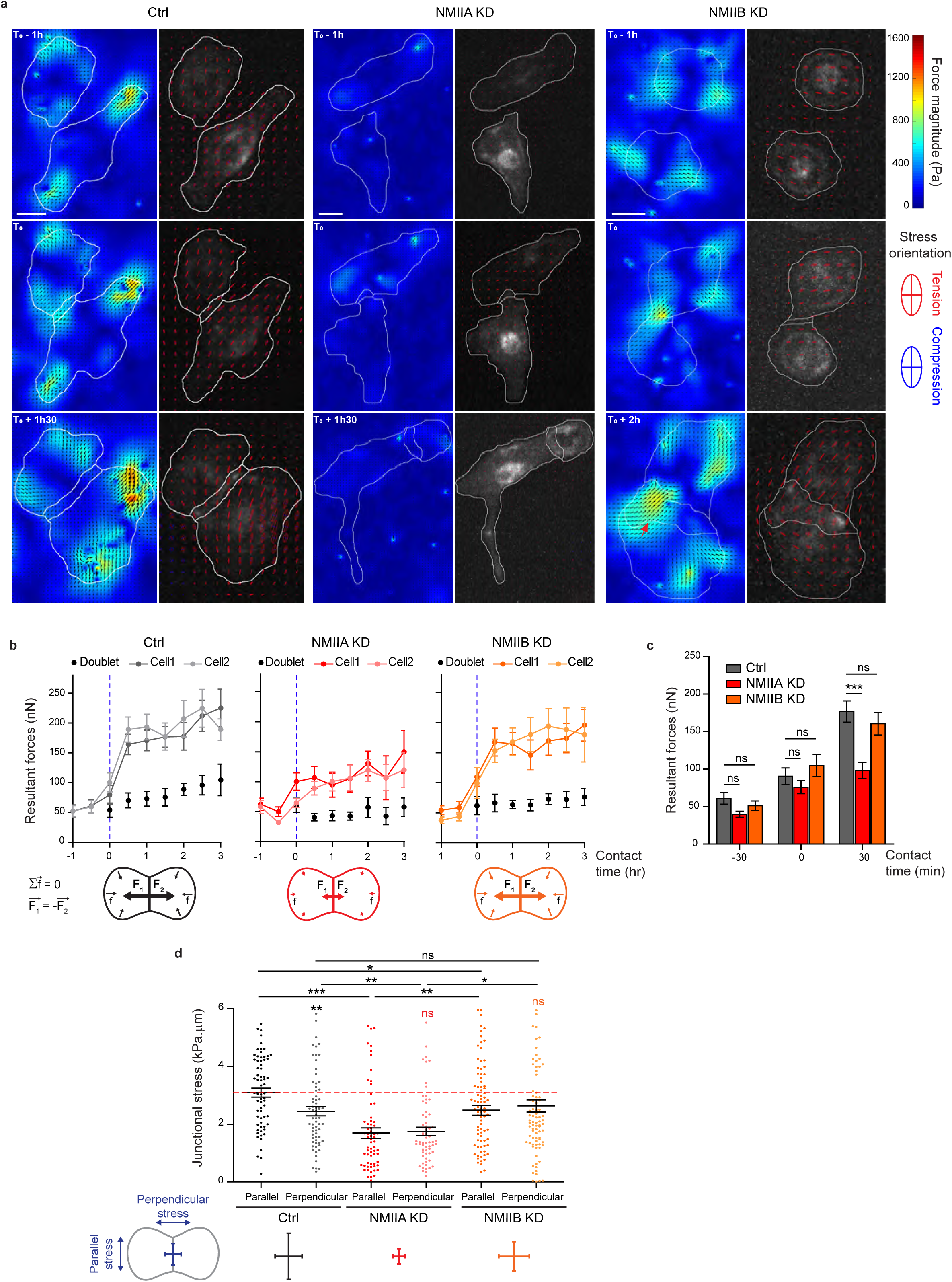
NMIIA and NMIIB are both required for establishment of proper inter-cellular stress. **a.** Heatmap with vectorial field of traction forces (left panels) and ellipse representation of intra-cellular stress (right panel, the two axes represent the direction and magnitude of the principal components of the stress tensor, positive values in red, negative values in blue) of inter-cellular stress (right panels) in Ctrl, NMIIA KD and NMIIB KD cell pairs before, during and after contact on fibronectin-coated PDMS deformable substrate (30 KPa). Cell contours are drawn in white. The red arrows indicate a hotspot of traction forces observed frequently in NMIIB KD cell doublets. Scale bar: 10 µm. **b.** Linear graphs representing the resultant forces of cell doublets and individual cells before, during and after contact in Ctrl, NMIIA KD and NMIIB KD. Data are represented as mean +/-SEM. **c.** The same data as in b were represented as bar graph with mean +/-SEM for statistical comparisons between Ctrl, NMIIA KD and NMIIB KD cells 30 minutes before, during and 30 minutes after contact. Bonferroni statistical tests were applied for p value. **d.** Scatter plots with mean +/-SEM representing inter-cellular stress in the junctional area in Ctrl, NMIIA KD and NMIIB KD cells within the first 3 hours of contact. The stress orientation was divided in the parallel and perpendicular components relative to the main axis of the junction. Mann-Whitney (for intra-group comparisons) and Kruskall-wallis statistical tests were applied for p value. **b-d**: n = 15, 20 and 18 cell doublets for Ctrl, NMIIA KD and NMIIB KD, respectively.

Using traction force measurement data, we then computed the intracellular stress in the cell doublets^85^ (Fig. 8a). The in-plane stress is represented by a tensor with three independent components: two components of normal stress denoting either tension (positive values) or compression (negative values) along the corresponding directions, and one component of shear stress, except in the basis of the tensor’s principle directions, where there is no shear stress. The ellipse representation in Figure 8a shows that the stress is highly anisotropic, and the cells are mostly under tension except for regions of very small compression associated to high tension in the other direction. The NMIIA KD cells show lower tension, consistent with the lower amount of traction forces they exert. We focused on the normal stress within the region of cell-cell junction, as AJs provide a mechanical link that drives transmission of forces between cells and thus organize inter-cellular stress^85,86^. We thus computed the perpendicular (σ_⊥_) and parallel (σ_//_) components of normal stress relative to the junction axis, which characterise the tension across and along the junction respectively. Within 30 minutes after contact formation, the junction was submitted to a rise of σ_⊥_ in all three cell lines, consistently with the emergence of a cell-cell tugging force (Supplementary Fig.5d,e). However, in Ctrl cells, the normal stress parallel to the junction σ_//_, remained higher than σ_⊥_ on average (Fig.8d). Strikingly, this was not the case in NMIIB KD and NMIIA KD cells that exhibited on average equal amounts of normal stress parallel and perpendicular to the junction, denoting a more isotropic distribution of junctional tension (Fig. 8d).

Altogether, these results show that NMIIA and NMIIB are both required for mechanical integrity of the junction. NMIIA is necessary for generation of a high junctional inter-cellular stress through production of tugging forces compensated by traction applied at cell-matrix adhesions. NMIIB, on the other hand, is necessary for the establishment of an anisotropic stress at the junction, sustaining high tension along the cell-cell interface.

## Discussion

Here, we explore for the first time the involvement of NMII isoforms during early steps of epithelial junction formation. We show that NMIIA and NMIIB associate with distinct pools of actin and cooperate to initiate the formation of epithelial AJ before the acquisition of the apico-basal polarization (See model, Supplementary Fig.6).

While NMIIA associated to actin bundles parallel to- and distant from the junction, NMIIB was sitting at junctional membranes in association with an Arp2/3-branched actin network, distinct from NMIIA-associated actin. The existence of two distinct actin networks at *adherens* junctions had already been observed in early junctions between hepatocytes^18^ and in endothelial cells where VE-cadherin was shown to colocalize with Arp2/3 complex-positive actin networks in-between distal actin-NMII bundles^65^. The localization of NMIIA is reminiscent of what has been observed previously in linear AJ of endothelial cells^65^. Strikingly, we show here an unexpected association of NMIIB with the Arp2/3-dependant branched actin that link the junctional membrane to NMIIA-associated perijunctional contractile actin bundles. Our data support a role of this isoform in organizing this branched network and its association to adhesion complexes on one side and perijunctional actin bundle on the other side. We believe that these are properties common to the early stage of AJ formation in many cell types^18,65^, that then mature to elaborate *zonulae adherens* in epithelial cells where both actin organizations, associated with their respective NMII isoforms, persist but become tightly packed to the junctional membrane^14,32^. Interestingly, in the absence of α-catenin KD, the localization of NMIIB was not restricted any more to junctional membranes in early epithelial junctions. Instead, NMIIB co-assembled with NMIIA on the same actin fibres, likely in heretotypic minifilaments, as observed in previous studies^36,61^, indicating that α-catenin is responsible for the junctional recruitment of NMIIB, as reinforced by a recent publication reporting NMIIB and α-catenin interaction in glioblastoma cells^73^.

These distinct localization patterns at early junctions are correlated to differential contributions of NMIIA and NMIIB in junction biogenesis. Upon contact formation, NMIIA KD cells were unable to elongate the junction and to sustain long-lived cell-cell contacts. They also lacked the capacity to produce traction forces on E-cadherin-coated substrates. Our observations thus identify NMIIA as the major isoform responsible for the NMII-dependent mechanical tugging force required for junction growth^29^. This was confirmed by traction force and stress analysis data revealing a decrease of the forces as well as a reduction of both parallel and perpendicular stresses at the junction for NMIIA KD cells. In contrast, NMIIB KD cells transmitted elevated tugging forces and maintained cell-cell contacts, but their junctions appeared enlarged and twisted with a lower parallel stress. These results are remarkable given that NMIIB was found to be expressed 100 times less than NMIIA in MDCK cells^35^. NMIIB was required for efficient E-cadherin clustering on E-cadherin substrates and for the connection of the contractile actin network to these clusters. NMIIB was also required for the proper organization and spatial restriction of the Arp2/3-dependent branched actin in the junctional area. Unexpectedly, NMIIB, and not NMIIA, was the main isoform responsible for the maintenance of α-catenin in an opened conformation.

Given that E-cadherin complexes have been shown to biochemically interact with both Arp2/3^16,77^ and NMIIB ^73^, one hypothesis could be that NMIIB and Arp2/3 are both recruited to E-cadherin/catenin complexes upon cell-cell contact initiation. NMIIB could thus serve as a cross-linker of the junctional actin network. Hence, the absence of NMIIB may induce a local softening of AJs which in turn leads to reduced extension of AJs and keeps α-catenin in a closed conformation. It is also in agreement with a previous study showing that Arp2/3-nucleated actin network at the *zonula adherens* regulates junctional tension and integrity^77^. NMIIB by associating both with cadherin-catenin complexes and the branched actin could somehow rigidify and regulate the thickness of this F-actin cushion sitting between the membrane and the contractile actin fibres associated to NMIIA. This could be achieved through the specific biochemical properties of NMIIB towards actin that provide it with the capacity to transmit tension within actin filaments at low energetic cost ^39,87,88^. Along this line, it is striking to note that we never observe in early AJ any sign of organization of NMIIB in minifilaments in the junctional area as observed for NMIIA in perijunctional actin bundles.

Inter-cellular stress is generated at cell-cell adhesions, although this remained poorly characterized^80,81^. Here, we evaluated the amount and the orientation of intercellular stress generated during junction biogenesis. Within one hour of cell-cell contact, an anisotropic inter-cellular stress appeared at the junction, with a preferential orientation parallel to the junction, favouring the elongation and the stability of the nascent junction. Both isoforms were required for proper establishment and orientation of this inter-cellular stress. NMIIA silencing had a global impact on the amount of inter-cellular stress generated, which was not surprising given its role on traction force production both at cell-matrix and cell-cell adhesions. On the other hand, NMIIB favoured the production of a higher parallel inter-cellular stress, probably by driving the crosslinking and stiffening of the junctional actin network that couples the perijunctional contractile actin to the plasma membrane.

In conclusion, we demonstrate here that both NMIIA and NMIIB contribute to the early steps of AJ biogenesis and are necessary for mechanical integrity of the junction, albeit implicated in very different aspects of adhesion complexes and actin pools organization. These findings open new avenues in the understanding of how distinct pools of actomyosin, associated to different myosin isoforms, built up and integrate mechanical forces to regulate *adherens* junction remodelling and intercellular stress in vertebrate cells in order to achieve large scale tissue remodelling during embryogenesis and tissue repair.

## Methods

### Antibodies and reagents

The following primary antibodies were used: rabbit anti-NMIIA polyclonal (Biolegend) or mouse anti-NMIIA monoclonal antibodies (Abcam, for co-immunostainings with anti-NMIIB antibodies); rabbit anti-β-catenin polyclonal (Sigma-Aldrich) or mouse anti-β-catenin monoclonal (BD Biosciences) antibodies; recombinant rabbit anti-paxillin monoclonal antibody (Abcam); mouse anti-GAPDH (ProteinTech), mouse anti-Arp3 (Sigma-Aldrich) and mouse anti-E-cadherin (BD Biosciences) antibodies; rabbit anti-α-catenin polyclonal (Sigma-Aldrich) and rabbit anti-NMIIB polyclonal (Biolegend) antibodies; rat anti-α18-catenin monoclonal antibody (generously provided by A. Nagafuchi (Kumamoto University, Japan)^89^. Alexa488-, Alexa568- and Alexa647-conjugated secondary antibodies were purchased from ThermoFisher, Alexa (488 or 555 or 647)-coupled phalloidins from Invitrogen and Hoechst 34580 from ThermoFisher. Horseradish peroxidase-coupled anti-mouse IgGs (Sigma-Aldrich) and anti-rabbit IgGs (Pierce) were used for immunoblotting. Mitomycin C and Y-27632 dihydrochloride were purchased from Sigma-Aldrich. The APP (Azido-Poly-lysine Poly (ethylene glycol)) and the BCN-RGD peptide (BCN: bicyclo[6.1.0]-nonyne, coupled to RGD: peptide sequence Arg-Gly-Asp) were prepared as previously described^90^.

### Cell culture

MDCK (ATCC CCL-34) and C2C12 (ATCC CRL-1772) cells originate from the American Type Culture Collection (ATCC). E-cadherin-GFP^91^ and α-catenin KD MDCK cell lines^92^ were kindly provided by W.J. Nelson (Stanford University, Palo Alto). Caco2BBE cells (ATCC HTB-37) were kindly provided by S. Robine (Institut Curie/CNRS, Paris). Cells were maintained at 37°C, 5% CO2 in DMEM (containing Glutamax, High Glucose and Pyruvate, Life Technologies) supplemented with 100 μg/mL Penicillin/Streptomycin (Life Technologies) and Foetal Bovine Serum (Life Technologies) at 10% for MDCK and C2C12 cells and at 20% for Caco2 cells. Ecadherin-GFP cells and α-catenin KD MDCK cells were maintained in media containing 5 μg/ml geneticin (Life Technologies).

### Generation of isoform-specific NMII knock-down MDCK cell lines

For generation of isoform-specific NMII Heavy chain knock-down cells, isoform-specific shRNA sequences, inserted in a back bone standard vector pLKO.1-puro, were designed and synthetized by Sigma-Aldrich technical services, based on the sequences of Canis lupus familiaris transcripts for MYH9 (NMIIA, transcript ID: ENSCAFT00000002643.3) and MYH10 (NMIIB, transcript ID: ENSCAFT00000027478). The sequences used were the following: TTGGAGCCATACAACAAATAC for NMIIA and TCGGGCAGCTCTACAAAGAAT for NMIIB.

As a control, the pLKO.1-puro non-mammalian shRNA Control Plasmid DNA was used (SHC002, Sigma-Aldrich). Two million Ecadherin-GFP MDCK cells were electroporated (Neon Transfection System Invitrogen) with 3-5 ug shRNA encoding plasmids in one pulse of 20 ms at 1650 V. Twenty four hours later, cells were put under selection pressure by adding puromycin (2.5 μg/ml) in media. After 10 days, single cells were sorted in 96 well plates by flow cytometry using Influx 500 sorter-analyzer (BD BioSciences) and clonal populations then selected based on NMII isoform expression levels by immunoblot and immunofluorescence. Control, NMIIA KD and NMIIB KD MDCK cells were maintained in media containing geneticin (5 μg/ml) and puromycin (2.5 μg/ml).

For simultaneous visualization of E-cadherin and centrosome, Ecadherin-GFP MDCK cells were transiently transfected with a plasmid driving the expression of RFP-Pericentrin (kindly provided M. Coppey, Institut Jacques Monod, Paris), using the protocol described above, one or two days before the experiment. m-Cherry cortactin plasmids (kindly provided by Alexis Gautreau, Biochemisty laboratory, Ecole polytechnique, France) were transfected in Control, NMIIA KD and NMIIB KD MDCK cells and the m-cherry expressing cell population was sorted by flow cytometry using Influx 500 sorter-analyzer (BD BioSciences).

### Western blotting

Confluent cells were lysed in 100 mM Tris pH 7.5,150 mM NaCl, 0.5% NP40, 0.5% triton-X100, 10% glycerol,1X protease inhibitor cocktail (Roche) and 1X phosphatase inhibitor (Phosphostop, Roche) for 20 minutes at 4°C. Insoluble debris were centrifuged for 15 minutes at 13000 g and supernatants were recovered. Protein concentration was quantified by Bradford assay (BioRad), SDS PAGE and electrotransfer were performed on 4-12% Bis-Tris gel (Novex) using mini gel tank and iBlot transfer systems (Invitrogen). Non-specific sites were blocked with 5% non-fat dry milk in PBS 0.1% Tween 20. Primary antibodies were diluted (1/1000) in PBS 0.1% Tween 20 and incubated overnight at 4ºC. After three washes in PBS 0.1% Tween 20, secondary HRP antibodies diluted in PBS 0.1% Tween 20 (1/10000) were incubated for 1 hour and washed 3 times with PBS 0.1% Tween 20. Immunocomplexes of interest were detected using Supersignal west femto maximum sensitivity substrate (ThermoFisher) and visualized with ChemiDoc chemoluminescence detection system (Biorad). Quantification of Western blots by densitometry was performed using the Gel analyzer plug in from Image J. GADPH was used as a loading control to normalize the quantification.

### Immunofluorescent staining

Cells were fixed with pre-warmed 4% formaldehyde in PBS for 15 min at RT and then washed 3 times with PBS, followed by permeabilization and blocking with 0.05% saponin/0.2% BSA in PBS for 15 minutes at RT. The primary antibodies diluted (1/100) in Saponin/BSA buffer were then incubated overnight at 4°C. After 3 washes in Saponin/BSA buffer, the samples were incubated with secondary antibodies and, when indicated, Alexa-coupled phalloidin, diluted at 1/200 in the same buffer for 1 hour at RT. The preparations were washed twice in Saponin/BSA buffer, once in PBS, and then mounted with the DAPI Fluoromount-G mounting media (Southern Biotech).

### Preparation of fibronectin-coated and cadherin-coated substrates

For fibronectin coating, glass coverslips were first cleaned by sonication in 70% ethanol and air dried. They were coated for 1 hour with 50 μg/mL human plasma fibronectin (Merck Millipore) diluted in PBS and washed three times with PBS.

The protocol for E-cadherin coating was inspired from a previous study by Lee and colleagues^93^. Briefly, the cleaned glass coverslips were silanized with 10% 3-aminopropyl triethoxysilane (APTES, Sigma-Aldrich) in 100% ethanol for 10 minutes at RT, washed once in 100% ethanol and dried at 80°C for 10 minutes. The surface was then functionalized by incubation for 1 hour with 2 mM EDC-HCl (Thermo Scientific) / 5 mM NHS (Sigma-Aldrich) and 1µg of recombinant human E-cadherin (R&D systems). Coverslips were then washed two times with PBS.

Cells were plated at very low density (typically 1 105 cells for a 32 mm diameter coverslip) on the coated coverslips in complete medium containing 10 μg/mL mitomycin C. After 1 hour incubation at 37°C, the preparations were washed twice with complete media and incubated 2-6 hours or overnight at 37°C before imaging or fixation, for cadherin coating and fibronectin coating, respectively.

### Preparation of switchable micro-patterns and imaging

Micropatterns were made as previously described with some modifications^94^. Briefly, air dried cleaned glass coverslips were activated with deep UV for 5 minutes, and coated for at least 1 hour with the repellent compound APP (0.1mg/ml in HEPES 10 mM pH7.4). After 3 washes with deionized water, the coverslips were exposed to deep UV for 7 minutes through a chrome photomask. The coverslips were then washed with deionized water three times, coated with 50 μg/mL human plasma fibronectin for 1 hour and washed twice with deionized water and once with PBS. When indicated, the coating was done with a 2:1 ratio of non-coupled:Cy3-coupled fibronectin prepared with Cy3 Mono-Reactive Dye Pack (GE Healthcare) as recommended by the manufacturer.

Cells were resuspended at 4.10² cells/mm² in medium containing 10 μg/mL mitomycin C and deposited on the patterned slide. After 1 hour of incubation at 37°C, cells were washed 3 times with fresh medium to remove mitomycin C and cells that remained in suspension. The cells that adhered on micro-patterns were left overnight in the incubator. The day after, confinement was released by addition of 20 μM BCN-RGD peptide diluted in DMEM media or, in case of live-imaging experiments, in Fluorobrite DMEM (Thermo Fisher) supplemented with 10% FBS and 1% Penicillin/Streptomycin. For ROCK inhibition experiments, 50 μM Y-27632 was added at the same time as BCN-RGD. Samples were then immediately imaged under a microscope or left in the incubator for 20 more hours and fixed as described above. When indicated for live-imaging experiments, nuclei were stained before adding BCN-RGD peptide by incubating the preparations with 5 μg/mL Hoechst 34580 in the medium for 20 minutes at 37°C followed by two washes with fresh media.

### Image acquisition and analysis

For live-microscopy experiments, the samples were placed in a chamber equilibrated at 37°C under 5% CO2 atmosphere. Images were acquired with a Yokogawa-Andor CSU-W1 Spinning Disk confocal mounted on an inversed motorised Leica DMI8 microscope and equipped with a sCMOS Orca-Flash 4 V2+ camera (Hamamatsu) and a 63 X oil immersion objective or a 20 X dry objective, with multi-positioning and a resolution of 0.5-3 μm z-stacks. Alternatively, the samples were imaged with an Olympus IX81 wide-field fluorescence microscope equipped with a Coolsnap HQ CCD camera and a 60X oil immersion objective or a 20 X dry objective. For some experiments, the Nikon Biostation IM-Q microscope was also used with 10X or 20X objective and multi-positioning.

For fixed samples, images were acquired with a Zeiss Apotome fluorescence microscope equipped with a 63 X oil immersion objective or with a Zeiss LSM 780 confocal microscope equipped with a 63 X oil immersion objective at a resolution of 0.3 μm z-stacks.

Image processing and analysis were done on Fiji software. Analysis of junction parameters (length, straightness and angle deviation) was done manually with Fiji software based on phase contrast and GFP-Ecadherin signal. Cell spreading, focal adhesions and α-catenin clustering were analyzed by thresholding the image and applying an “Analyze particles” which gives the number of objects and its area. To calculate the ratio of α-cat to α18-cat intensities, the mean grey intensity value for the two channels were measured within the manually-defined junction. Tracking of single cells on fibronectin was done using the Manual Tracking plugin.

### Traction force microscopy

Soft silicone elastomer substrates for TFM (Traction force microscopy) were prepared as described previously with some modifications^95^. Cy 52-276 A and Cy 52-276 B silicone elastomer components (Dow corning) were mixed in a 5:5 (elastic modulus ∼15kPa for E-cadherin-coating) or a 5:6 ratio (elastic modulus ∼30kPa for fibronectin-coating). 0.08 g of elastomer was deposited on 32 mm glass coverslips and allowed to spread progressively. The substrate was silanized with 10 % (3-aminopropyl triethoxysilane (APTES, Sigma) in 100% ethanol for 10 minutes at RT, washed once in 100 % ethanol and dried at 80°C for 10 minutes. The surface was coated for 10 minutes at RT with carboxylated red fluorescent beads (100nm, Invitrogen) diluted at 2-3/1000 in deionized water. After washing with deionized water, the surface was finally functionalized with protein (fibronectin or E-cadherin) as described above. Seeded cells together with fluorescent beads were imaged either on an Olympus-CSU-W1 Spinning Disk confocal microscope with a 10 X dry objective and 3 μm z stacks or on an Olympus-IX81 wide field inverted fluorescence microscope with a 20 X dry objective for 2 to 24 hours, at a frequency of 1 frame every 10 min, at 37°C under 5% CO2 atmosphere. At the end of the acquisition, 100-200 μL of 10% SDS was added in the media to detach cells and image a reference frame. For force calculation, matPIV was used to analyse the displacement vectors of the beads, which were further translated into forces using the FTTC plugin in ImageJ. The vector quiver plots and heat map of magnitude force was plotted using Matlab. Mean (resp. resultant) forces exerted by cells and doublets were obtained by computing the average of the magnitude (resp. the vectorial sum) of traction forces within manually defined masks. For the analysis of tractions forces below cell-cell junctions, the junction masks and corresponding midline were first manually defined based on the E-cadherin-GFP pictures. Then, the midline was used to define the average orientation of the junction, and all force vectors within the junction mask were projected onto the directions parallel and perpendicular to this orientation. The mask was divided in four quarters along this mean orientation. The “junction centre parallel (resp. perpendicular) force” is defined as the averaged absolute value of the parallel (resp. perpendicular) component of traction forces in the two central quarters of the mask, while the “junction periphery parallel (resp. perpendicular) force” is the averaged absolute value of the parallel (resp. perpendicular) component of traction forces in the two outermost quarters.

T_(parallel / perpendicular)^(center/periphery)=〈|T_(parallel / perpendicular) |〉_(center / periphery)

### Calculation of inter-cellular stress

Computing the junctional stress components σ= andσ_⊥_, respectively parallel and perpendicular to the cell junction (Fig. 8d), required both the determination of the cell junction location and the estimation of the inter-cellular stress tensor. The cell junction domain was defined as the overlap between two masks representing the area covered by each cell in the doublet. Given the stress tensor, the parallel and perpendicular stress components were obtained by rotation from the cartesian basis. As exemplified in Fig. 8a, we found in most cases that the cell junction domain was roughly straight: the mean orientation of the cell junction domain determined the rotation angle. We checked that following the cell junction contour did not significantly modify our estimates. Finally, each junctional stress component was spatially-averaged over the cell junction domain.

Intercellular stress was estimated by Bayesian inversion^96^, with a dimensionless regularization parameter Λ= 10^5^ (see ^8^ for details). The spatial domain for stress estimation was for each image the smallest rectangle encompassing the cell doublet. For simplicity, we implemented free stress boundary conditions on the straight boundaries of the rectangular domain, instead of following the cell doublet boundaries. As a consequence, the stress estimation was qualitative, but sufficed to evaluate differences between conditions. Note that height variations within the cell doublet were also neglected in the estimation of the 2D inter-cellular stress field.

### SIM microscopy

Super-resolution structured-illumination microscopy was performed on a Zeiss Elyra PS.1 microscope with a 63 X objective (Plan Apo 1.4NA oil immersion) and an additional optovar lens 1.6 X. Cells grown on 0.17 mm high-performance Zeiss coverslips were fixed and prepared for immunostaining, then with DAPI Fluoromount-G mounting media (Southern Biotech). Laser lines 488 nm, 561 nm and 641 nm were directed into the microscope, passing through a diffraction grating. For 3D SIM imaging, the diffraction grating was rotated along 3 directions (angles 120o) and translated (five lateral positions) throughout the acquisition. Typically, 20-30 slices of 110 nm were acquired for each cell corresponding to an imaging height of 2-3 μm. The fluorescence signal was detected with an EMCCD camera (iXon-885, Andor, 1004×1002, pixel size 8 μm, QE=65%). Processed SIM images were aligned via an affine transformation matrix of predefined values obtained using 100 nm multicolor Tetraspeck fluorescent microspheres (Thermo Fisher Scientific).

### Data display and statistics

Images were mounted using Photoshop and Illustrator. Graphs and statistical tests were done using GraphPad prism software.

## Acknowledgements

This work was supported by the European Research Council (Grant No. CoG-617233), LABEX “Who Am I?,” and the Agence Nationale de la Recherche “POLCAM” (Grant No. ANR-17-CE13-0013). We acknowledge the ImagoSeine core facility of the Institut Jacques Monod, member of IBiSA and France-BioImaging (ANR-10-INBS-04) infrastructures. We thank Orestis (ImagoSeine core facility, IJM) for technical assistance with SIM experiments. We thank Sree Vaishnavi and Gianluca Grenci (Micro fabrication Core Facility of Mechnabiology Institute, National University of Singapore) for the fabrication of chrome photomask. We thank A. Nagafuchi for α18-catenin antibody, W.J. Nelson and S. Robine for providing cells, M. Coppey for RFP-Pericentrin plasmid and M. Piel (IPGG, Curie Institute) for providing original APP and BCN-RGD compounds. We thank Delphine Delacour and Shreyansh Jain for useful scientific discussions and critical reading of the manuscript.

## Author contributions

R.M.M, B.L. and M.H.L. conceived the project. R.M.M and B.L. supervised the project. M.L.H, G.S and T.D performed experiments. M.L.H, G.S., R.M.M and B.L. designed experiments and analyzed data. J.A., V.C and P.M analyzed traction force microscopy data and calculated inter-cellular stress. D.W and J.v.H designed and performed the production of APP and BCN-RGD compounds. M.L.H, G.S., R.M.M, B.L. J.A., V.C and P.M wrote the manuscript.

---

**Supplementary figure 1:**
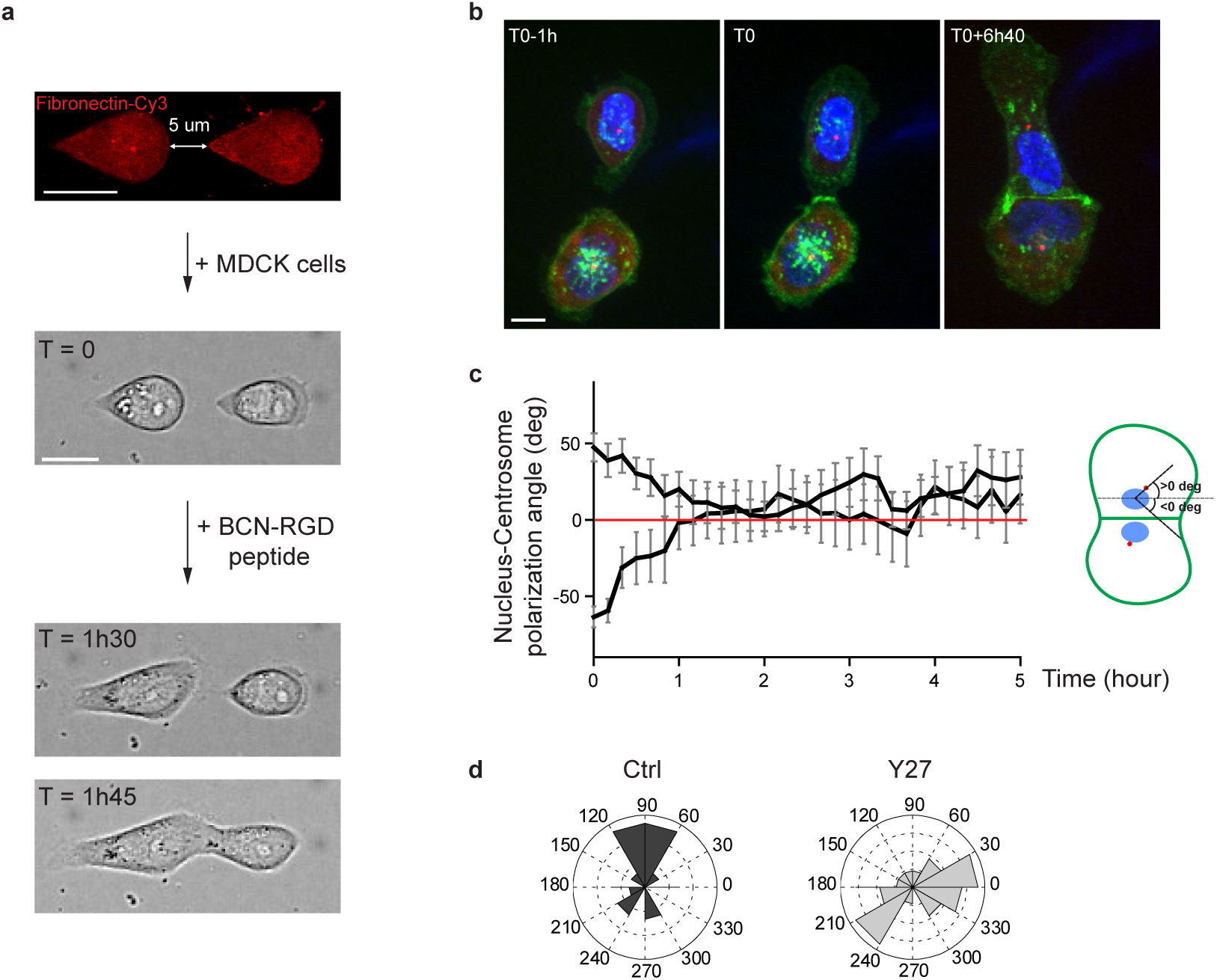
Reversal of nucleus-centrosome polarity axis after cell-cell contact. **a.**Sequential steps for controlled initiation and visualization of junction biogenesis. The two cells are initially confined on a pair of fibronectin-coated 5 µm-away patterns (T=0). When desired, the cell confinement is released by addition of BCN-RGD peptide, inducing cell spreading and kissing within a few hours. Scale Bar: 10 µm. **b.** Spinning disk image sequence of GFP-E-cadherin and RFP-Pericentrin of doubled transfected MDCK cells pre-stained with Hoechst. Scale bar: 10 µm. **c.** Plots of nucleus-centrosome axis polarization angle relative to junction axis after cell-cell contact. Data are represented as mean +/-SEM. n = 19 doublets from three independent experiments. **d.** Distribution of nucleus-centrosome axis polarization angles relative to junction axis after cell-cell contact in Ctrl or Y27-treated MDCK cells at the time when junctions reach their maximal length. n = 15 and 21 cells from three and two independent experiments respectively.

**Supplementary figure 2:**
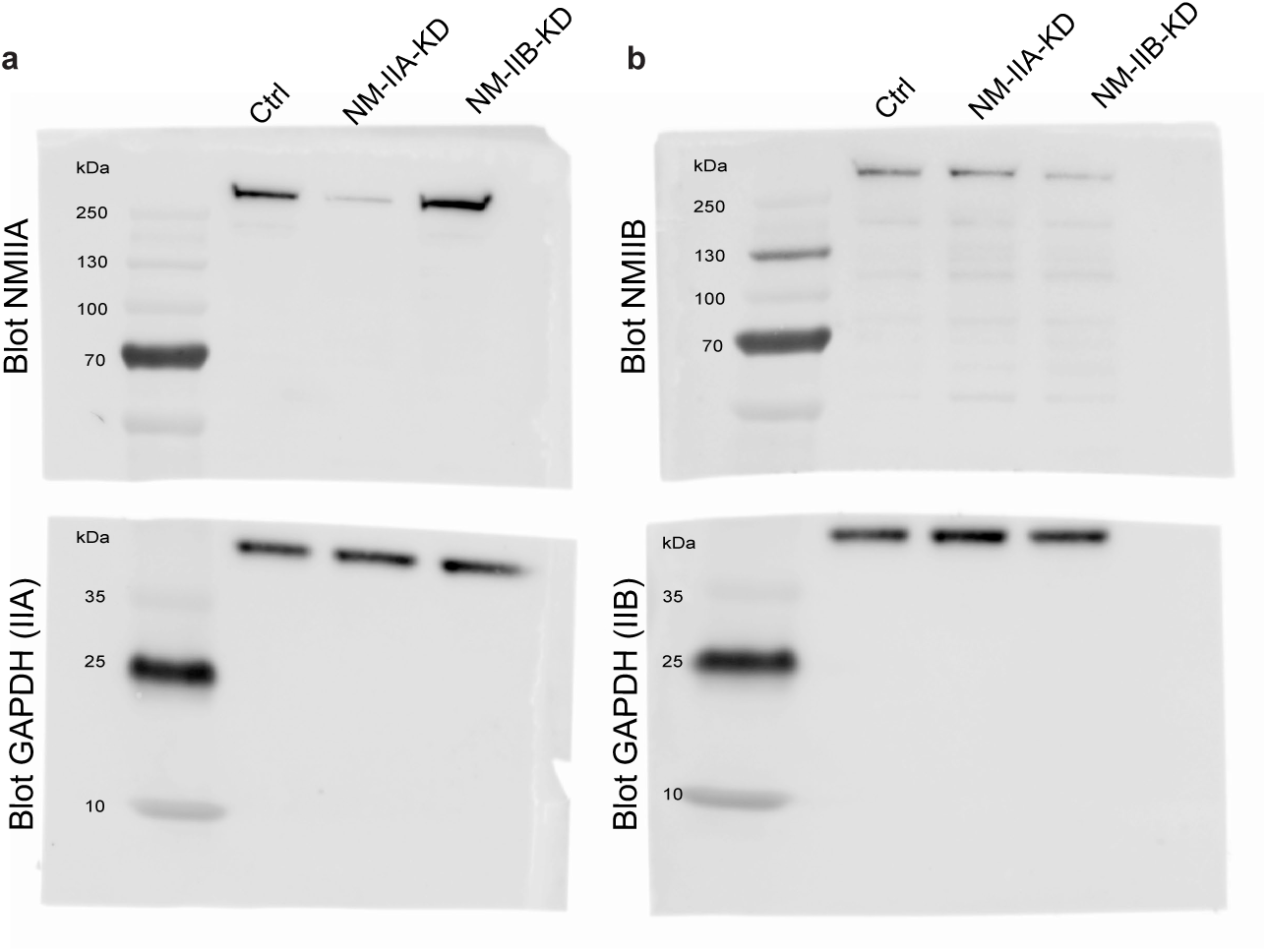
Isoform-specific NMII Knock-down in MDCK cells. **a.b.** Original uncropped Immunoblots presented in Figure 2a (a) and 2b (b).

**Supplementary figure 3:**
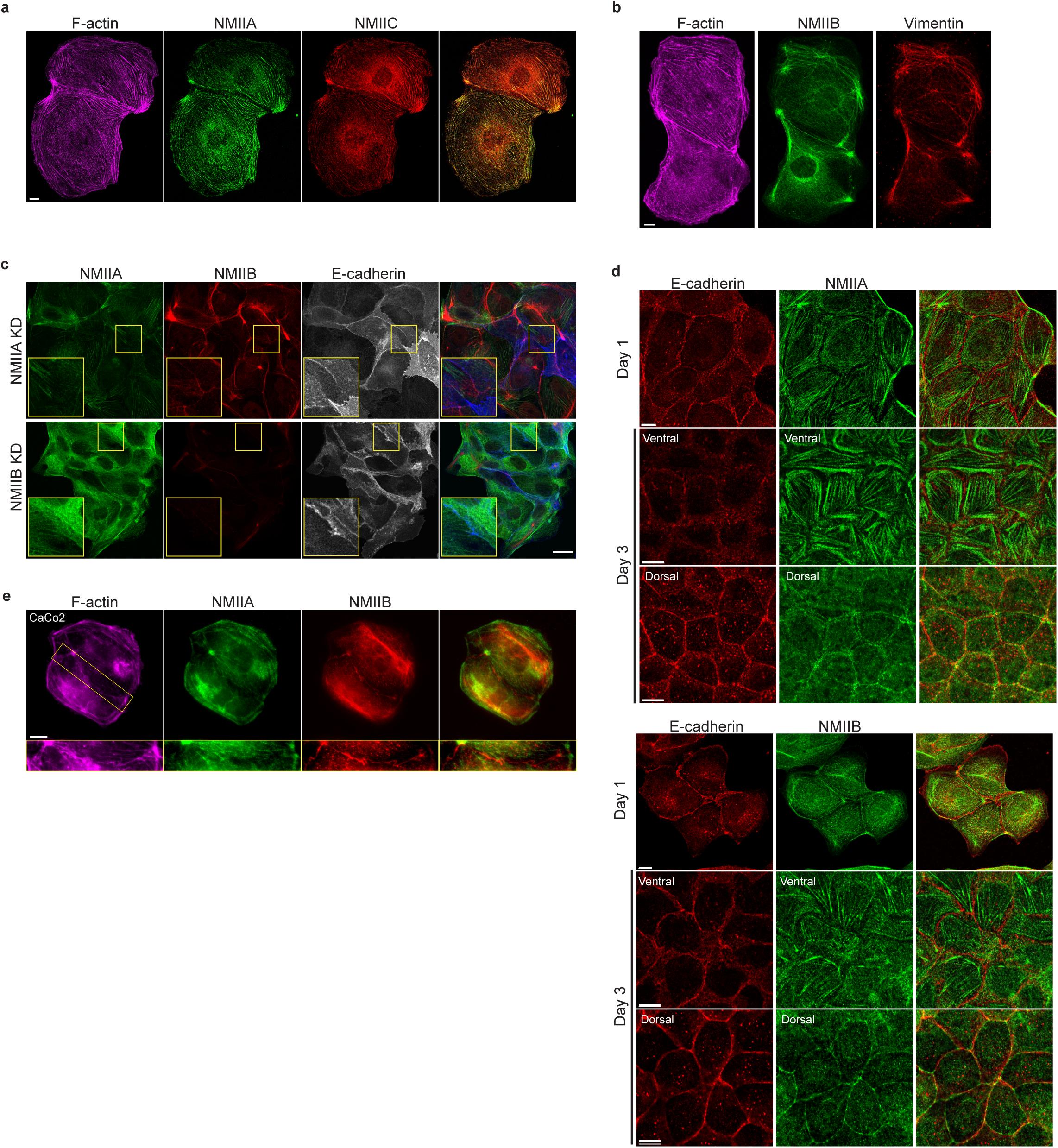
NMIIB, but not NMIIA, localizes to early epithelial AJs. **a.b.** Representative confocal images of MDCK cell doublets fixed 20h after BCN-RGD addition and stained for F-actin, NMIIA and NMIIC (a) or F-actin, NMIIB and Vimentin (b) as indicated. Scale bar: 10µm. **c.** Representative confocal images of NMIIA KD and NMIIB KD MDCK cells plated at low density on fibronectin, fixed after 12 hours and immuno-stained for NMIIA, NMIIB and E-cadherin. Scale bar: 20µm. **d.** Representative confocal images of WT MDCK cells plated on fibronectin-coated glass for 1 or 3 days and immuno-stained for E-cadherin and NMIIA (left panel) or NMIIB (right panel). Scale bar: 10µm. **e.** Representative epifluorescent images of Caco-2 cells plated at low density on fibronectin, fixed after 12 hours and stained for indicated proteins. Scale bar: 10µm.

**Supplementary figure 4:**
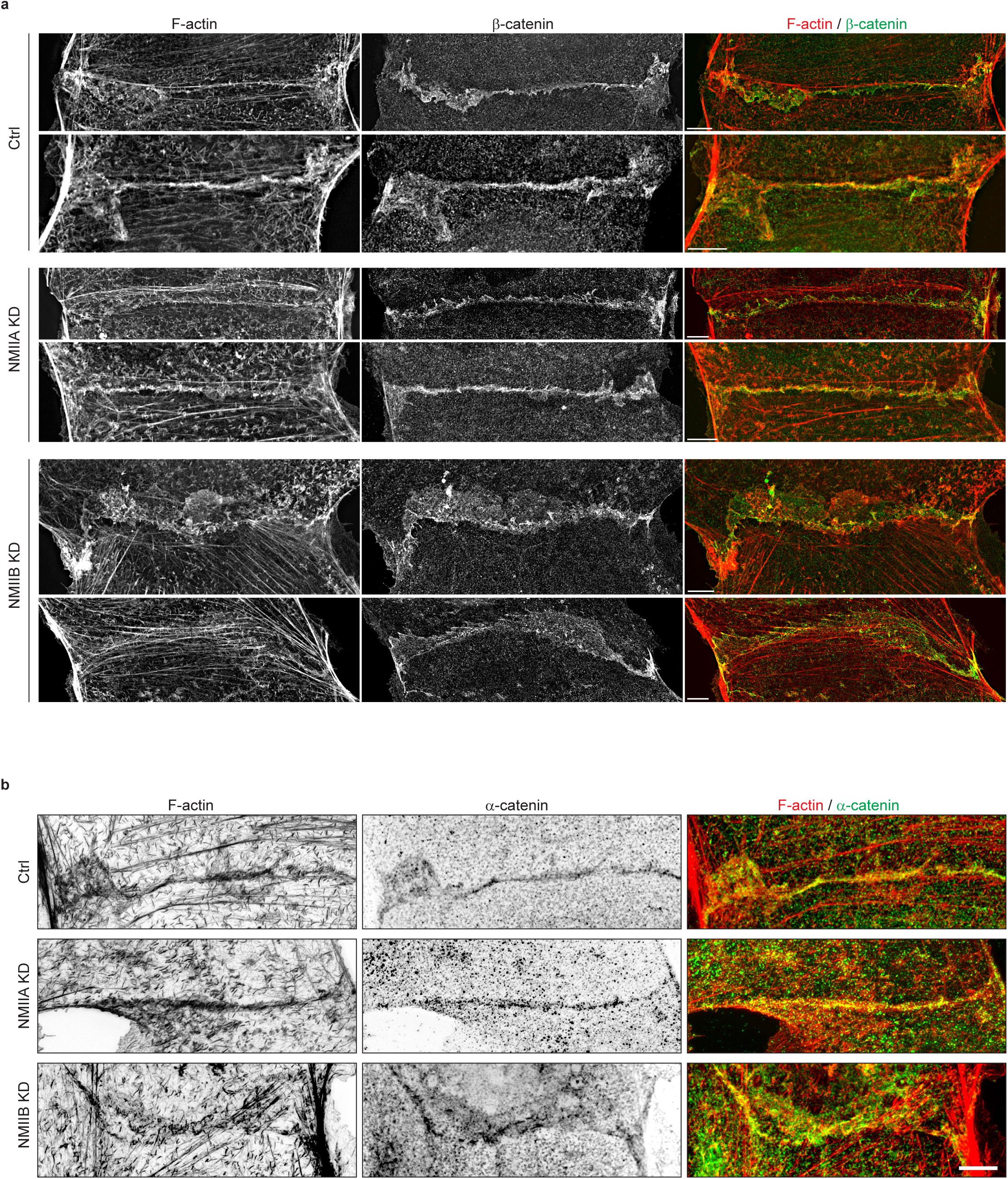
NMIIB supports junctional actin organization. Related to Figure 5a: other examples of junctional actin organization in Ctrl, NMIIA KD and NMIIB KD cells. **a.b.** SIM (Structured Illumination Microscopy) images of junctional area from Ctrl, NMIIA KD and NMIIB KD cells fixed 20h after addition of BCN-RGD and stained for F-actin and β-catenin (a) or α-catenin (b). Scale bar: 5 µm.

**Supplementary figure 5:**
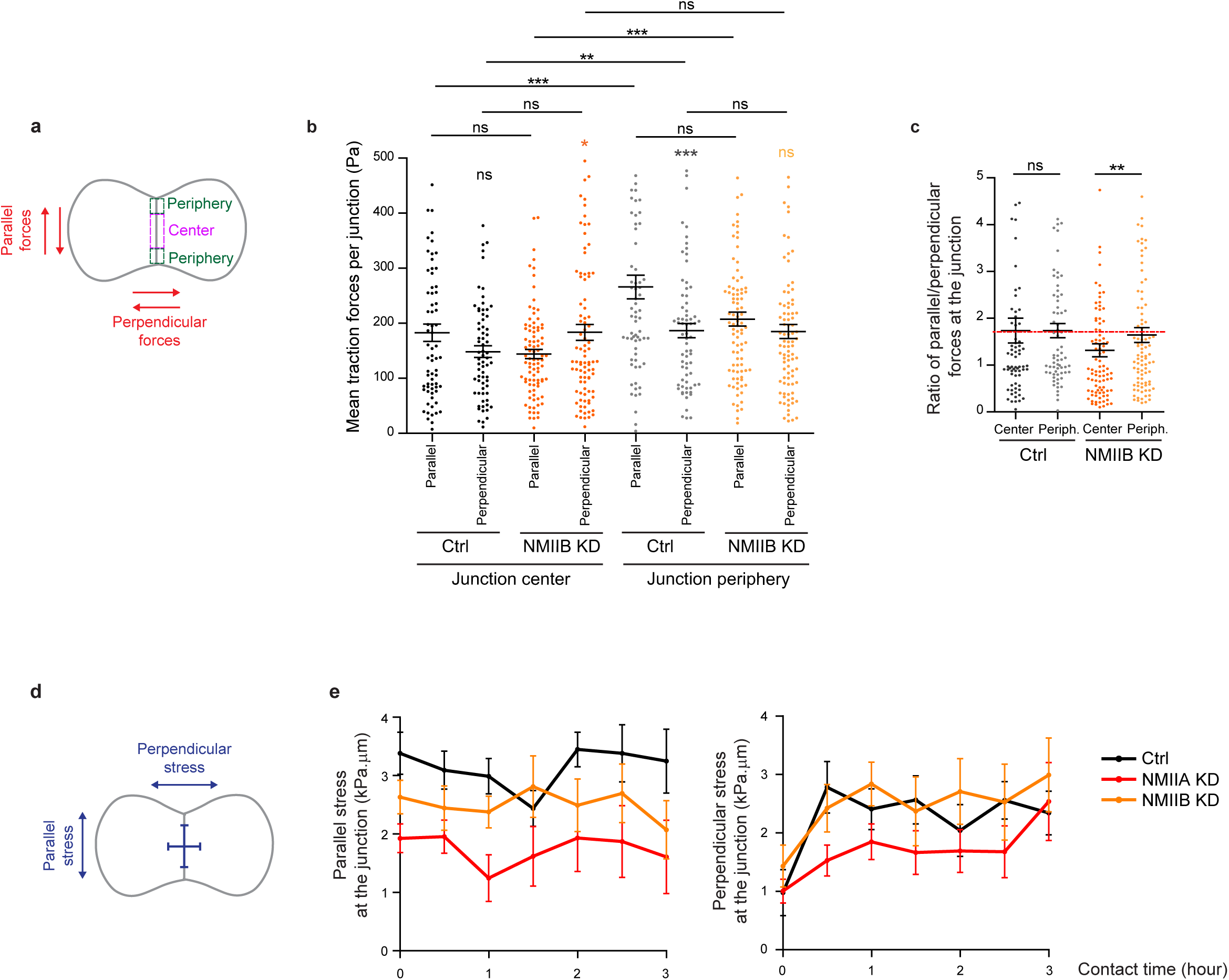
NMIIA and NMIIB are both required for establishment of proper inter-cellular stress. **a.**Scheme depicting the junction subdomains and the orientation of traction forces relative to the junction axis quantified in (b) and (c). **b.c.** Scatter plots with mean +/-SEM representing the orientation of mean traction forces (b) and the ratio of parallel/perpendicular traction forces (c) in subdomains of the junction in Ctrl and NMIIB KD cells within the first 3 hours of contact. Mann-Whitney (for intra-group comparisons) and Kruskall-wallis statistical tests were applied for p value. n = 15 and 18 cell doublets respectively. **d.** Scheme depicting the orientation of inter-cellular stress relative to the junction axis quantified in (e). **e.** Linear graph of parallel (left panel) and perpendicular (right panel) inter-cellular stress at the junction within the first 3 hours after contact in Ctrl, NMIIA KD and NMIIB KD cells. Data are represented as mean +/-SEM. n= 15, 20 and 18 cell doublets for Ctrl, NMIIA KD and NMIIB KD respectively.

**Supplementary figure 6:**
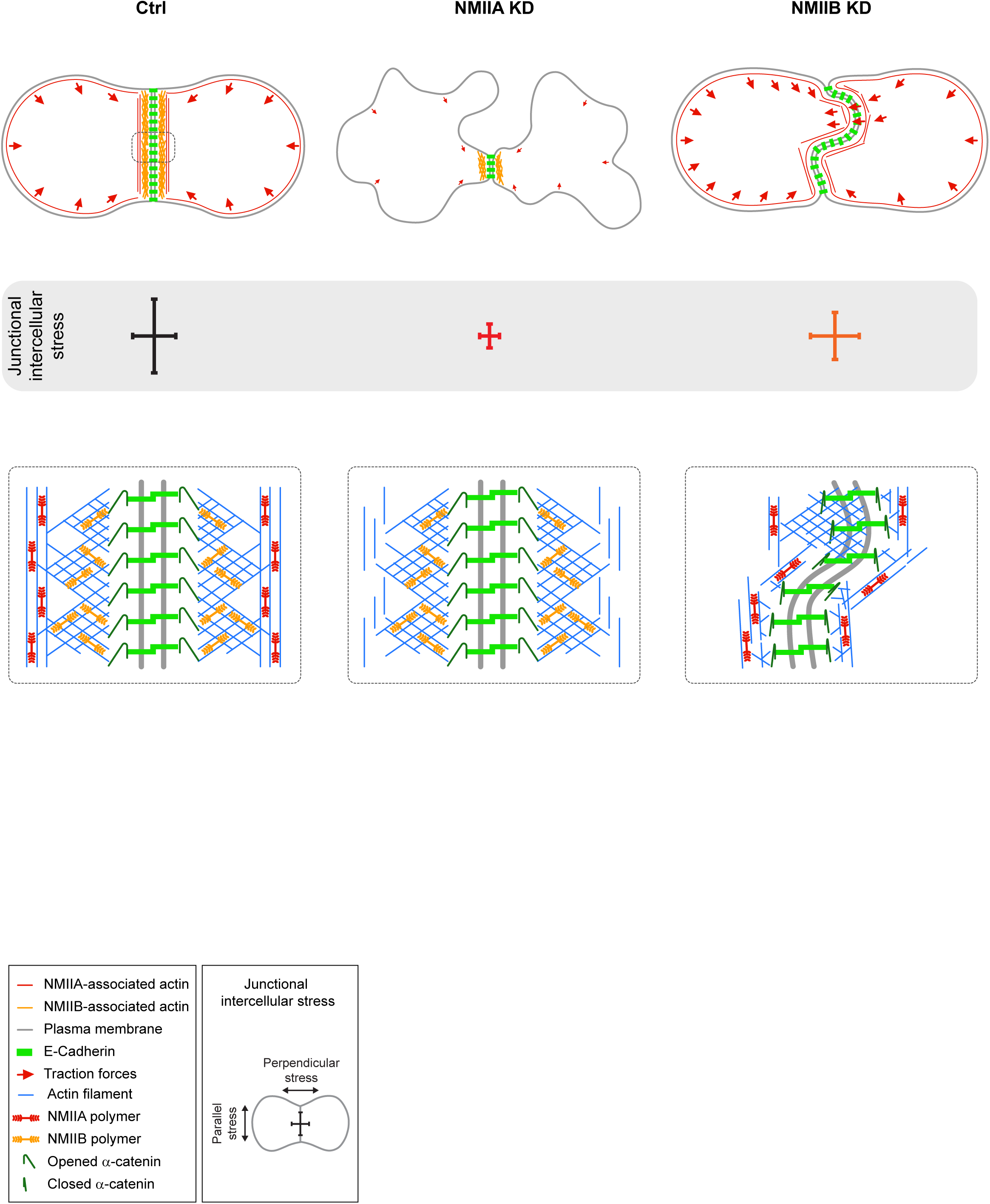
Proposed model for the role of NMIIA and NMIIB during junction biogenesis. Upper panels: summary of the phenotypes observed in Ctrl, NMIIA KD and NMIIB KD cells during junction biogenesis. Lower panels: proposed molecular organization of early junctions in all three cell lines, based on the results obtained in this study and previous ones (see Discussion part). Ctrl cells establish stable and straight junctions with traction forces mainly peripheral, generating an anisotropic intercellular stress preferentially parallel to the junction. NMIIB associates to- and organizes the junctional branched actin meshwork, and favours the opening of α-catenin molecules. NMIIA, which provides mechanical tugging force, sits on distant perijunctional actin bundles parallel to the junction. Actin cables parallel to the cortex and ventral stress fibers (not represented here for clarity of the figure). NMIIA KD cells fail to keep contact for long times, exhibit shorter junctions, weak traction forces and weak intercellular stress. Perijunctional actin bundles are smaller and disorganized. The junctional NMIIB-actin meshwork still supports α-catenin opened conformation but it does not prevent the junction from disassembly. NMIIB KD cells establish persistent but wavy junctions from which lamellipodial extensions and traction force hotspots arise. There is no preferential orientation of intercellular stress. The junctional branched actin meshwork is disorganized, which probably prevents α-catenin opening and induces the formation of lamellipodial extensions. The anchoring of perijunctional actin bundles to the junction is perturbed, despite the presence of NMIIA.

## Supplementary Video legends

**Supplementary Video 1: Dynamic of junction formation on reversible micropatterns**

Spinning disk movie showing contact formation between two MDCK cells expressing GFP-E-cadherin and stained with Hoechst. Scale bar: 10 µm.

**Supplementary Video 2: Dynamic of junction formation in Y27-treated cells**

Spinning disk movie of MDCK cells expressing GFP-E-cadherin, stained with Hoechst and treated with 50 µM Y27. Scale bar: 10 µm.

**Supplementary Video 3: Dynamic of junction formation in Ctrl, NMIIA KD and NMIIB KD cells**

Epi-fluorescent movies of Ctrl, NMIIA KD and NMIIB KD MDCK cells expressing GFP-E-cadherin. Scale bar: 10 µm.

